# Role of Phage–Antibiotic Combinations in Reducing ESBL-Producing and Carbapenem-Resistant *Escherichia coli*

**DOI:** 10.1101/2024.06.28.601134

**Authors:** Md Shamsuzzaman, Shukho Kim, Jungmin Kim

## Abstract

The emergence of extended-spectrum *β*-lactamase (ESBL)-producing *E. coli* and carbapenem-resistant *E. coli* (CREC) poses a significant global health concern. Here, we isolated and characterized two novel phages and studied their effectiveness with antibiotics against ESBL-producing *E. coli* and CREC. The isolated phages, EC.W1-9 and EC.W15-4, belonged to the *Podoviridae* and *Myoviridae* families, respectively. They are safe for bacterial control as they do not contain integrase or toxin-coding genes. The phage combination considerably enhanced lytic ability, effectively lysing 61.7% of the 60 *E. coli* isolates, compared to lysis in the 41.6% –55% range by individual phages. Phages EC.W1-9 and EC.W15-4 combined demonstrated 100% susceptibility against different *E. coli* sequence types, including ST73, ST648, ST2311, ST405, ST7962, ST131, ST13003, and ST167. Additionally, studies showed synergy between antibiotics and phage combinations against ESBL-producing *E.coli*, with susceptibility of 73.3% and 54% for CREC. The combined treatment of isolated phages and antibiotics significantly increased survival rates in BALB/c mice exposed to various ST types of ESBL-producing *E. coli* and CREC, including ST131, ST648, and ST410. Survival rates against KBN7288 (ST131) increased by approximately 75% and 50% compared to individual phages EC.W1-9 and EC.W15-4, respectively. When phages and antibiotics were combined, survival rates against *E. coli* isolates KBN5617 (ST410), KBN6241 (ST410), and KBN4004 (ST648) ranged from 75% – 100%. Finally, this study highlights the importance of phage and phage-antibiotic combinations to prepare phages for killing different ST types of ESBL-producing *E. coli* and CREC isolates.

**IMPORTANCE:** When combined with antibiotics, phage therapy shows promise in fighting multidrug-resistant bacteria. However, antagonism between phages and antibiotics has been reported. This research isolates and characterizes two novel phages, EC.W1-9 and EC.W15-4, from the *Podoviridae* and *Myoviridae* families, respectively, and evaluates their effectiveness against ESBL-producing *E. coli* and CREC. These phages, lacking integrase or toxin-coding genes, showed significant promise in bacterial control. Combined phage treatment lysed 61.7% of *E*. *coli* isolates, outperforming individual phages. The phage combination showed 100% susceptibility against different *E. coli* sequence types. Additionally, the synergy between phages and antibiotics increased susceptibility rates to 73.3% for ESBL-producing *E. coli* and 54% for CREC. In BALB/c mice, combined treatments significantly improved survival rates against various *E. coli* isolates. Finally. this study emphasizes the potential of phage and phage-antibiotic combinations in targeting various ST types of ESBL-producing *E. coli* and CREC.

## INTRODUCTION

Antimicrobial resistance (AMR) is a significant global public health threat, with over 700,000 people dying annually from infections caused by AMR. Experts predict that 2050 annual deaths due to AMR will reach 10 million, costing $100 trillion (1, 2). The prevalence of extended-spectrum *β*-lactamases (ESBLs) significantly contributes to the current antibiotic resistance crisis. Even before the advent of penicillin in medicine, *E. coli* demonstrated resistance to *β*-lactam antibiotics, underscoring the importance of understanding *β*-lactamases (3). *β*-lactamases developed resistance to various antibiotics, including penicillin, third-generation cephalosporins (such as ceftazidime, cefotaxime, and ceftriaxone), and the monobactam aztreonam (1, 4). Previously, TEM and SHV type ESBLs were the most prevalent ESBL families (5). Today, CTX-M-type enzymes are the most common ESBL type, with the CTX-M-15 variant dominating globally, followed by CTX-M-14 and CTX-M-27 emerging in a particular region (6, 7). ESBL-encoding genes are frequently found on plasmids and within transposons or insertion sequences, facilitating their easy spread. Moreover, the global population of ESBL-producing *E. coli* is dominated by a highly virulent and successful clone of the ST131 strain (8). ESBL-producing organisms frequently exhibit co-resistance to various antibiotic classes, which limits therapeutic options (9). Recent studies have demonstrated fluoroquinolone resistance mediated by co-transfer of the *qnrA* determinant on ESBL-producing plasmids, highlighting the broad antibiotic resistance extending to multiple antibiotic classes (10). Despite their resistance, clavulanic acid can effectively inhibit ESBLs, yet these enzymes confer multi-resistance to antibiotics and related oxyimino *β* -lactams, posing significant challenges to treatment (9, 11).

In 2017, the World Health Organization (WHO) identified CREC as a critical, highest-priority organism, emphasizing the urgent necessity for new antibiotics to address its impact (12). CREC is known to cause severe infections such as intra-abdominal infections, pneumonia, urinary tract infections, and device-associated infections (13). The high levels of antibiotic resistance and frequent transmission between humans and animals make CREC infections a significant public health concern (1). In addition, CREC is also a significant food safety concern (14). The abundance of antibiotic resistance and virulence genes in CREC strains reduces the effectiveness of traditional antibiotics and complicates patient management (1). Moreover, CREC has evolved into a complex and resistant bacterium, posing challenges to treatment strategies (15, 16). One study observed the prevalence of various sequence types (STs) of CREC isolates in 874 *E. coli* isolates carrying *bla*_NDM-5_ collected from 13 European Union or European economic area countries over 2012–2022 (17). Another study highlighted the growing concern of high-risk CREC isolates, specifically *E. coli* ST131, circulating within the intensive care units in China (18). Furthermore, researchers suggest compelling evidence of CREC transmission among humans, animals, and the environment (19, 20).

Phage therapy, using phage to treat bacterial infections, has gained renewed interest as a potential solution in the fight against antimicrobial resistance (21). Phages, which are highly abundant and diverse viruses in the biosphere, exhibit a specific and efficient bactericidal effect. They can serve as antibiotic alternatives or be combined with antibiotics for synergistic effects (22). Clinical approaches for phage therapy have been reported in several countries, such as the United States, Georgia, Poland, and Russia (23, 24). Scientific evidence supporting the benefits of phage therapy for humans and animals has steadily increased in recent decades (25). The limited availability of phages underscores the urgent need to isolate safe, highly lytic, and well-characterized phages for phage therapy (26). However, bacteria growing to resist phage is a big problem, even though it is a natural process that helps keep different kinds of germs around, which is vital for the environment’s health (27). The emergence of phage-resistant bacterial strains is almost unavoidable, as frequently reported in *in-vitro* studies and clinical trials (28, 29).

Phage therapy necessitates strategies to control the emergence of phage-resistant bacterial strains. This can be accomplished through the utilization of phage cocktails, which consist of multiple therapeutic phages, or by combining phages with antibiotics (30). Phage cocktails are employed to target various structural sites and metabolic activities of a bacterium, reducing the likelihood of simultaneous mutations affecting all receptors (31). One study has shown that a phage cocktail was utilized to treat *P. aeruginosa* infection in mice, and no phage-resistant bacterial mutants were observed despite being identified during *in vitro* experiments (32). While phage cocktails have shown promise in preventing the emergence of resistant bacteria, it is important to note that phage resistance has been observed in the case of *E, coli* strains, and there is limited knowledge about the elimination of the resistant bacteria *in vivo* (33). Previous studies have reported that phage-resistant mutants are less frequently observed in animal models or during clinical trials, but there are exceptions (28, 29, 32). This study evaluated the efficacy of phage combination (EC.W1-9 and EC.W15-4) and phage-antibiotics (synergistic activity) in eliminating the different ST types of ESBL*-*producing *E. coli* and CREC isolates *in vitro* and *in vivo*. Furthermore, the study also evaluated the combination of phage antibiotic synergistic against the phage and antibiotic-resistant clinical isolate CREC ST410.

## RESULTS

### Host ranges of novel *E. coli* phages and their combination

Based on the spot test results, phages EC.W1-9, EC.W15-4, and their combination exhibited high lytic activity against 14 different *E.coli* ST types, including 60 ESBL-producing and CREC isolates. The lytic activity of the isolates was observed at 43.3% (26/60), 55.0% (33/60), and 63.3% (38/60), respectively (Table 1 & Fig. S1A). Moreover, the lytic activity against ESBL-producing *E.coli* was about 53.8 % (14/26), 61.5% (16/26), and 73.0% (19/26) (Fig. S1B), Where the lytic activity against CREC was found to be 37.5% (18/48), 50.0% (24/48) and 56.2% (27/48) (Fig. S1C). Additionally, phage EC.W1-9 and EC.W15-4 combination demonstrated 100% susceptibility against 9 *E. coli* sequence types (n=14), including ST131(n=18), ST648 (n=4), ST7962 (n=2), ST73 (n=1), ST2311(n=1), ST405 (n=1), ST1487(n=1), ST13003 (n=1), and ST167 (n=1) (Fig. S1D). However, the susceptibility of ST410 to the phage combination was approximately 34.6% (9/26). While single phage EC.W1-9 and EC.W15-4 showed susceptibilities of 11.5% (3/26) and 26.9% (7/26), respectively. The efficiency of plating (EOP) values of phage EC.W1-9 against *E. coli* KBN 7288 (ST131), KBN4004 (ST648), and KBN6241 (ST410) compared to reference *E. coli* ATCC25922 are approximately 1.9×10^8^, 2.9×10^4^, and 1.462×10^9^ respectively. For phage EC.W15-4, EOP values are about 4.8×10^5^ and 6.8×10^8^ for *E. coli* KBN7288 and KBN4004 (Table S1).

**Table 1.**
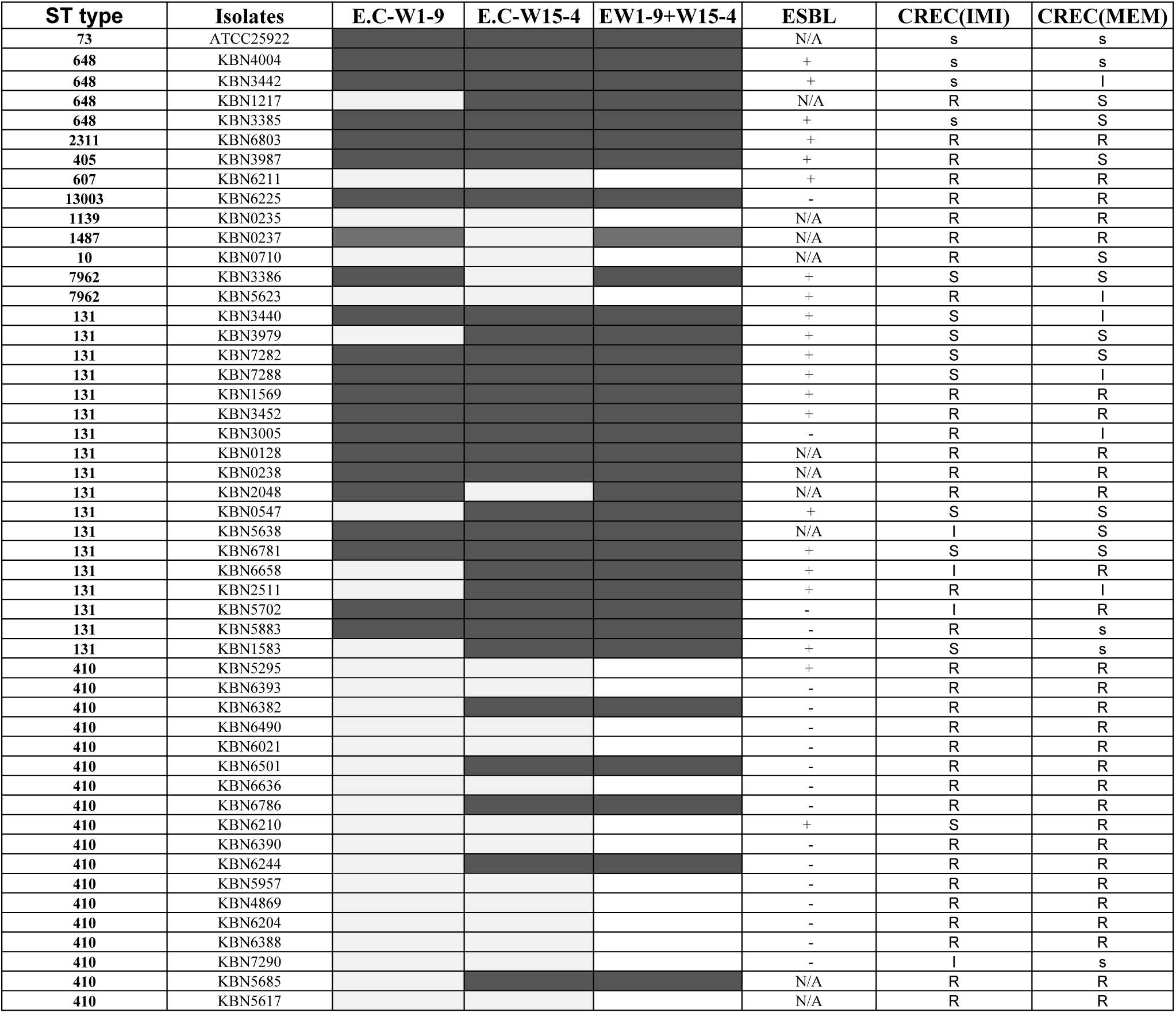
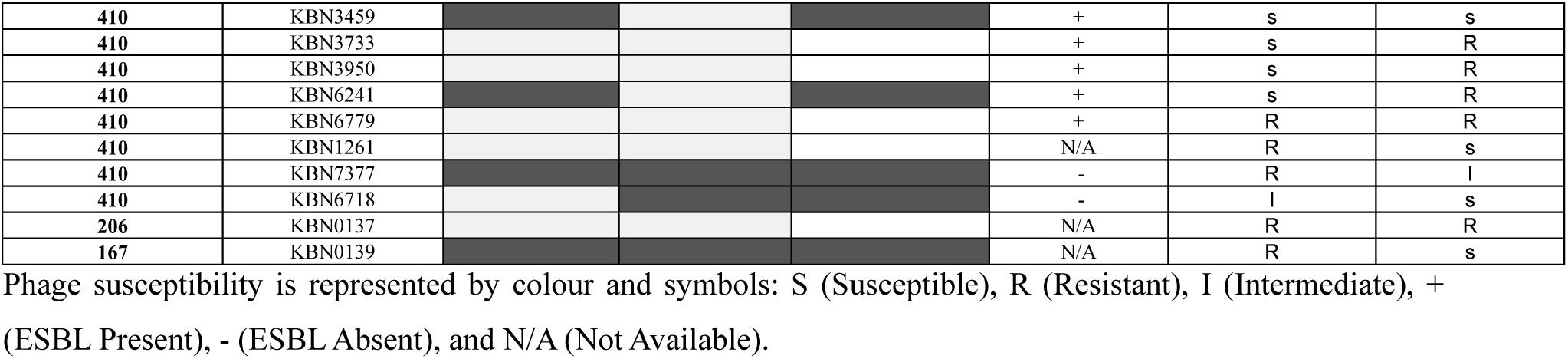
Host range of phages EC.W1-9 and EC.W15-4 and their combination against 14 different ST type of ESBL-producing *E.coli* and CREC.

### Antibiotic resistance profile of clinically isolated ESBL-producing and CREC isolates

The study analyzed 60 *E. coli* isolates and found that all were multidrug-resistant, with 80.4% being CREC isolates (34). Additionally, 43.33% of the isolates carried extended *β* -lactamase (ESBL). These isolates displayed significant resistance to various antibiotics, including cephalosporins (cefazolin (100%, n=41), ceftazidime (95.0% n=60), cefoxitin (85.3% n=41), ceftriaxone (100%, n=9), cefotaxime (97.5%, n=40), and cefepime (93.3% n=60)), aminopenicillins (amoxicillin (95.5%, n=45) and ampicillin (100% n=38)), monobactams (aztreonam (94.8%, n=58)), carbapenems (imipenem (77.0%, n=60) and meropenem (66.6%, n=60)), fluoroquinolones (ciprofloxacin (91.1%, n=58) and levofloxacin (94.4, n=18)), aminoglycosides (amikacin(26.3%, n=57), tobramycin (66.6%, n=3), and gentamicin (62.0%, n=58)), penicillins (piperacillin (30%, n=10)), sulfonamides (trimethoprim (87.1%, n=39)), tetracycline (100%, n=1), and oxazolidinone (linezolid 100%, n=1) (Table S2).

### Biological and morphological characterization

The study investigated the lytic activity, growth curve, thermal stability, and pH stability of phages EC.W1-9 and EC.W15-4. The phages exhibited better lytic activity against *E. coli* ATCC25922 at MOIs of 10 and 0.1 after 24 h of incubation at 37°C (Fig. 1). Within 24 h, the OD600nm values of the positive control increased continuously from 0.05 to 1.25, while that of the negative control remained unchanged. Treatment with phages significantly inhibited *E. coli* ATCC25922 growth at all MOIs within the first 8-9 h. However, the OD600nm values gradually rose after 8 h of phage treatment. After 24 h, there was a significant difference in the OD600nm values compared to the positive control and a considerable difference in MOI values. The OD600nm values continued to rise gradually until reaching 24 h. Although the bacterial count for all MOIs exceeded 10^8^ CFU/ml at 24 h, there was a significant difference between MOI 10 and 0.001 compared to the positive control. Interestingly, MOI 0.001 exhibited better lytic activity in the first 12 h than other MOI values. However, after 24 h, MOI 10 showed higher lytic activity. These findings demonstrate that these phages can significantly inhibit bacterial growth at all MOIs, with the highest bactericidal activity observed at MOI 10 and 0.1 after 24 h.

**Figure 1.**
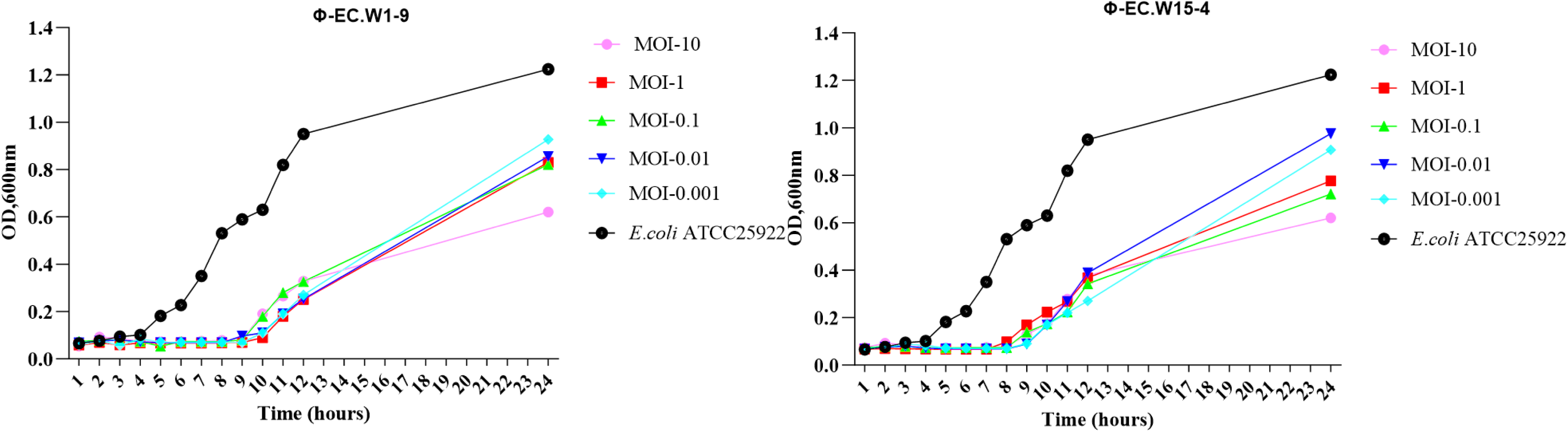
*In vitro* bacterial lytic activities of phages at various MOIs. Killing curves of *E. coli* (ATCC25922) by two phages at MOIs of 10,01, 0.1,0.01 0.001, and for 24h. Each point represents the means ± SD of three replicate experiments.

According to the phage adsorption assay, 90 > % of phages could adsorb to *E. coli* ATCC25922 within 10 min (Fig. 2A-B). In one-step growth curve analysis, the latent period of phages is approximately 15 min, followed by a rapid release of virus particles. The final titer reached a range of 10^7.5^-10^8^ PFU/mL after a burst period of 35 min, with a burst size of about 110-85 phage particles per cell (PFUs/cell) (Fig. 2C-D). The phages showed thermal stability between 4°C and 50°C for 60 min (optimal 4 to 50°C). Phage titers significantly decreased at 70°C and were completely inactivated at 80°C (Fig. 3A-B). They also showed better stability in different pH media at pH-3 to 10 within 4 h (optimal pH at 6 to 8) (Fig. 3C-D). After 18 h of incubation at 37°C on a double-layer agar plate, clear plaques with a diameter of about 1-2 mm were formed. Transmission electron microscopy (TEM) images showed that EC.W1-9 had an icosahedral head measuring 60 ± 5 nm in diameter and a noncontractile tail measuring about 05 ± 1 nm in length. The EC.W15-4 phage had a head size of 84 ± 3 nm and a contracted tail size of 100 ± 4 nm (34).

**Figure 2.**
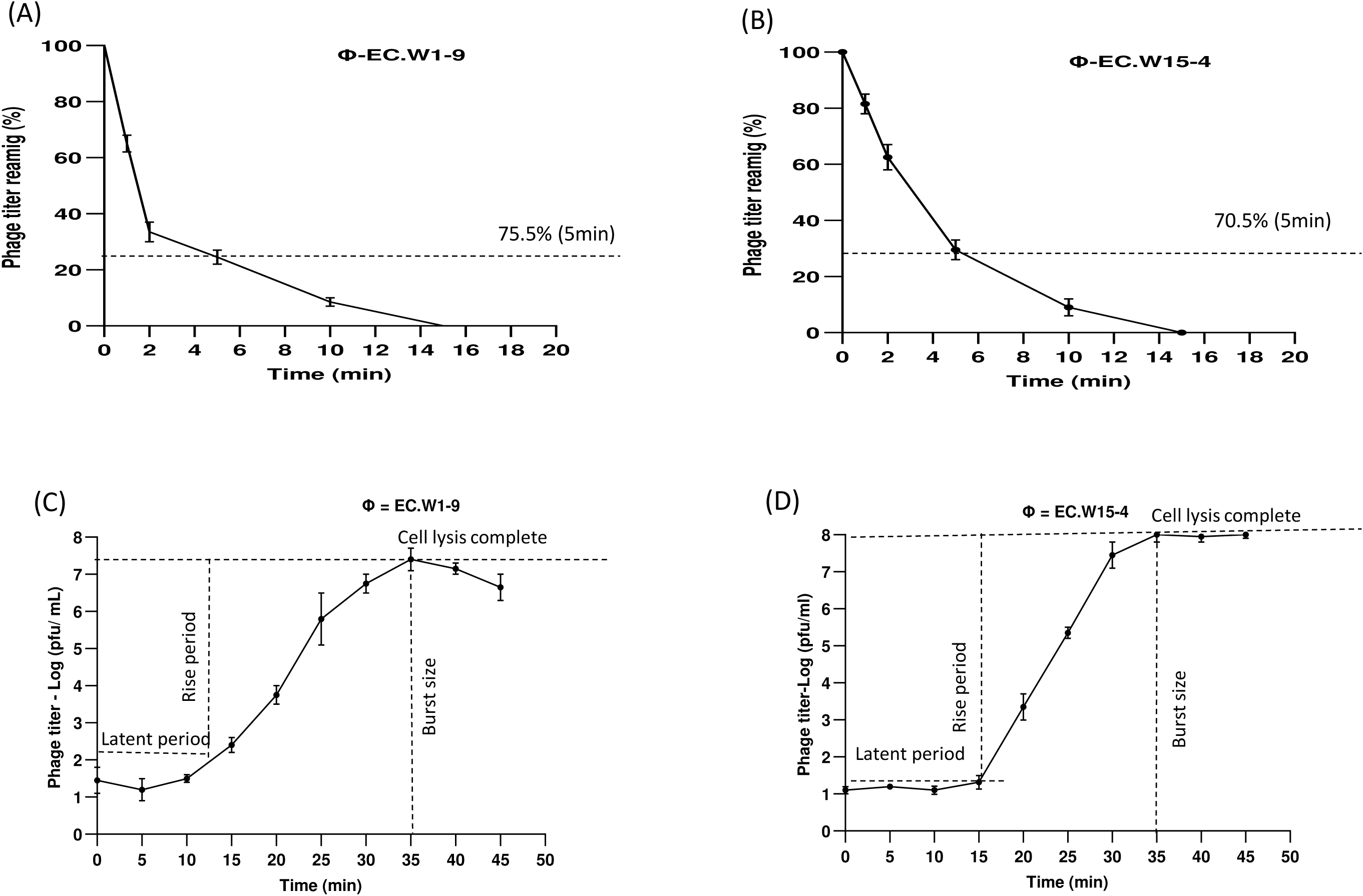
Adsorption rate and burst size of phages EC.W1-9 and EC.W15-4 to *E. coli* ATCC25922. 2(A-B) Adsorption assay measuring the percentage of remaining free phages over 15 minutes at MOI of 0.0001 and 2(C-D) One-step growth curve showing the latent period and burst size of two *E. coli* bacteriophages in BHI medium at MOI of 0.0001. Values represent means ± standard deviations from the duplication of each treatment.

**Figure 3.**
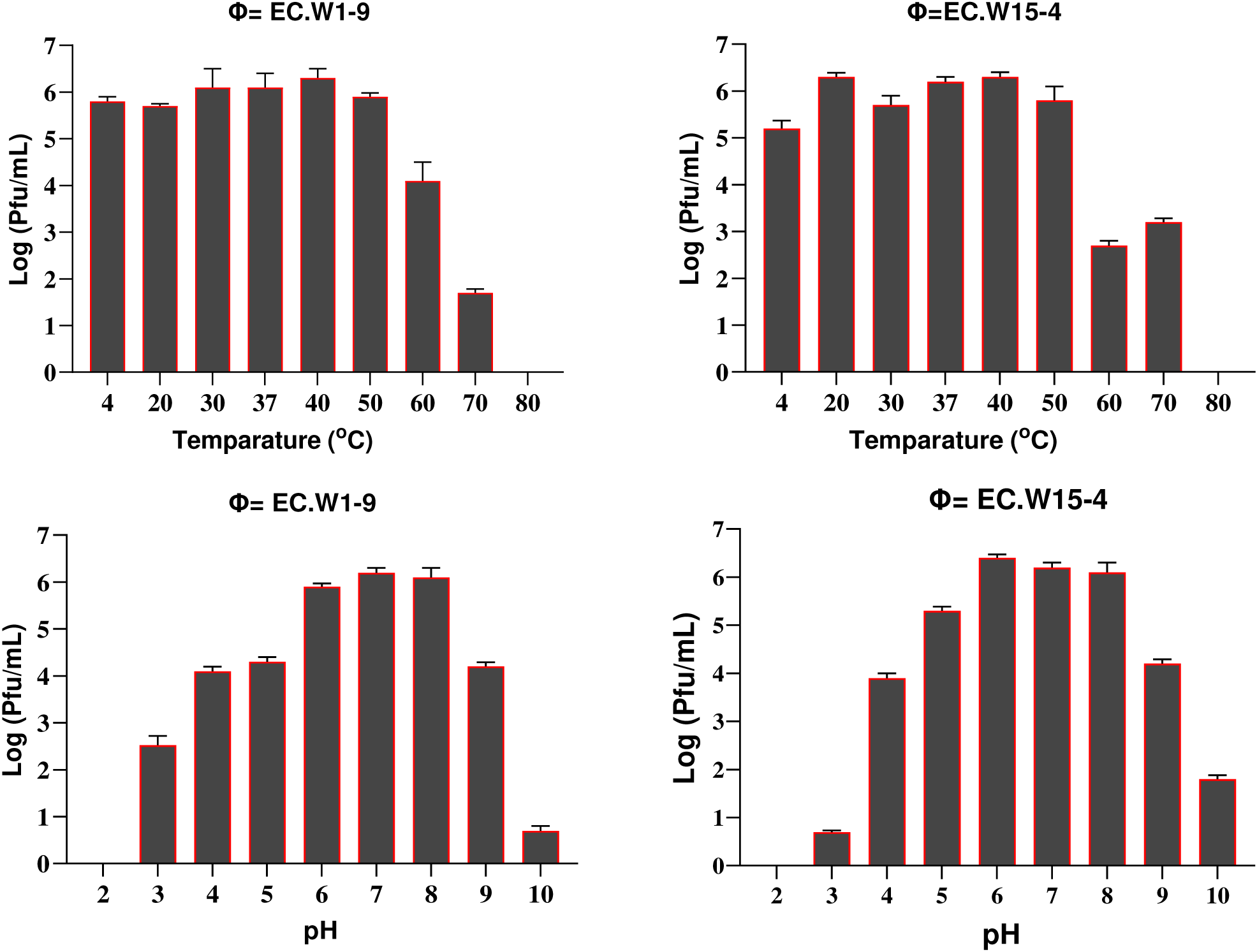
Stability of two novel *E. coli* bacteriophages. 3(A-B) Effect of temperature on phage stability, with phages treated at 4, 20, 30, 37, 40, 50, 60, 70, and 80°C for one hour, and surviving phage titers calculated. 3(C-D) pH stability of phages, with concentrations (∼10^8^ PFU/mL) incubated at pH 2 to 10 at 37°C for four hours. Each point represents the means ± SD of three replicate experiments.

### Genomic feature of novel phages EC.W1-9 and EC.W15-4

The whole-genome sequence analysis indicated that phage’s EC.W1-9 and EC.W15-4 had a circular double-stranded DNA structure, with respective genome sizes of 40,616 bp, and 152,184 bp. Additionally, their G+C content was found to be 45.8%, and 36.1% respectively (Table 2). Phylogenetic analysis using VIPtree revealed that phage EC.W1-9 is closely related to Escherichia phage phiV10 (NC_007804.2) and Escherichia phage TL-2011b (NC_019445.1), while EC.W15-4 is closely related to Escherichia phage vB_EcoM_G9062 (NC_054920.1) and Escherichia phage vB_EcoM_fFiEco06 (NC_054914.1). Phage EC.W1-9 belongs to the novel *Uetakevirus* genus in the *Podoviridae* family, and Phage EC.15-4 belongs to the *Tequatrovirus* genus in the *Myoviridae* family (Fig. 4.) In addition to the novel isolates EC.W1-9 and EC.W15-4, along with their highest DNA sequence similarities, the closest phages displayed a high degree of similarity. More than 95% of their genomes were conserved when aligned using the MAUVE alignment tool. This suggests that these species are closely related and share recent common ancestors. The MAUVE alignment revealed small, unaligned regions in the genome sequences, marked by white lines within blocks. These variable and conserved areas suggest functional differences and unique characteristics among the phages (Fig. 5).

**Figure 4.**
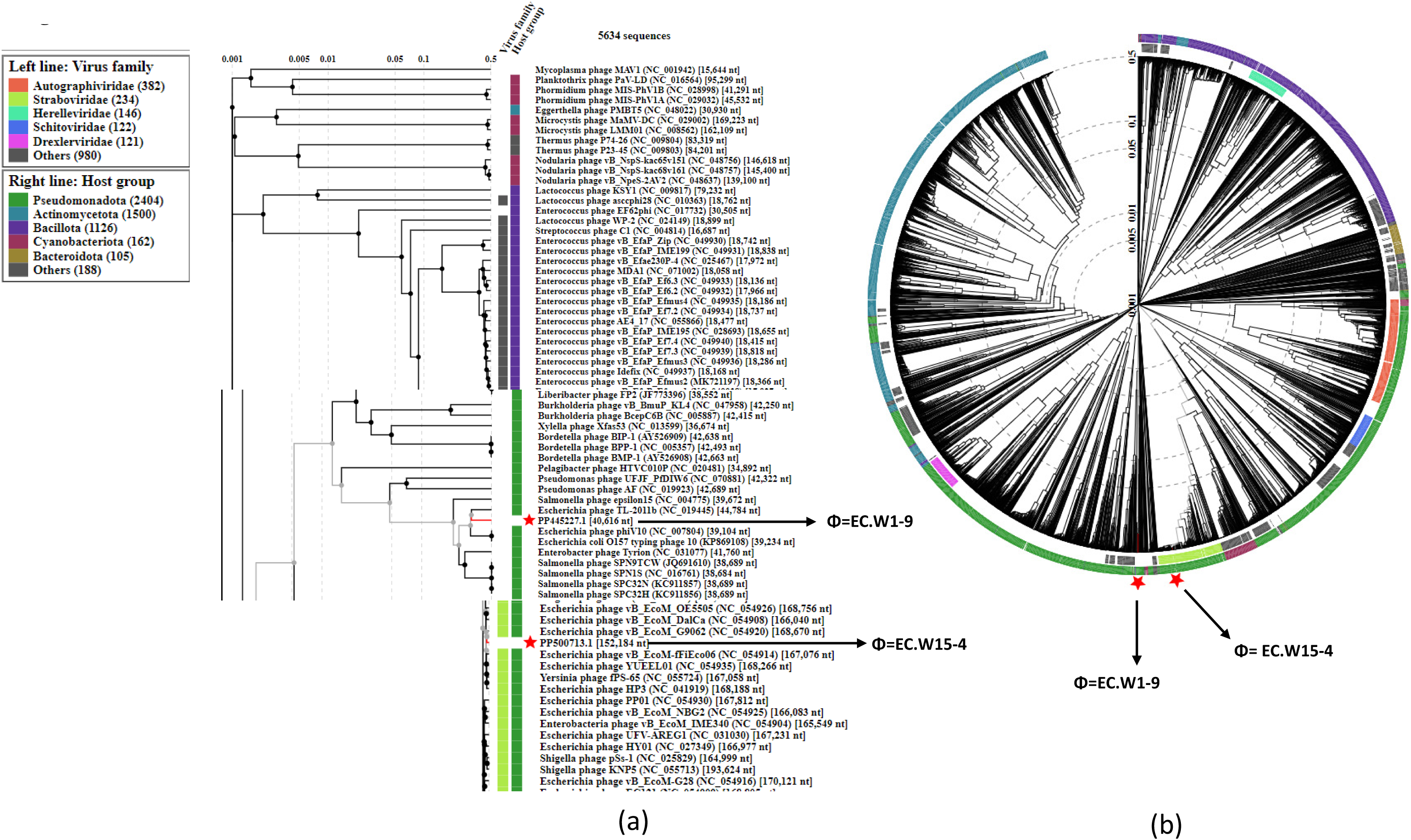
Position of two *E. coli* phages in the Phage Proteomic Tree. (a) The rectangular presentation shows the closest related phages to our isolates, indicated by a red asterisk. (b) Circular proteomic tree of prokaryotic dsDNA viruses, colour-coded by virus families and host taxonomic groups.

**Figure 5.**
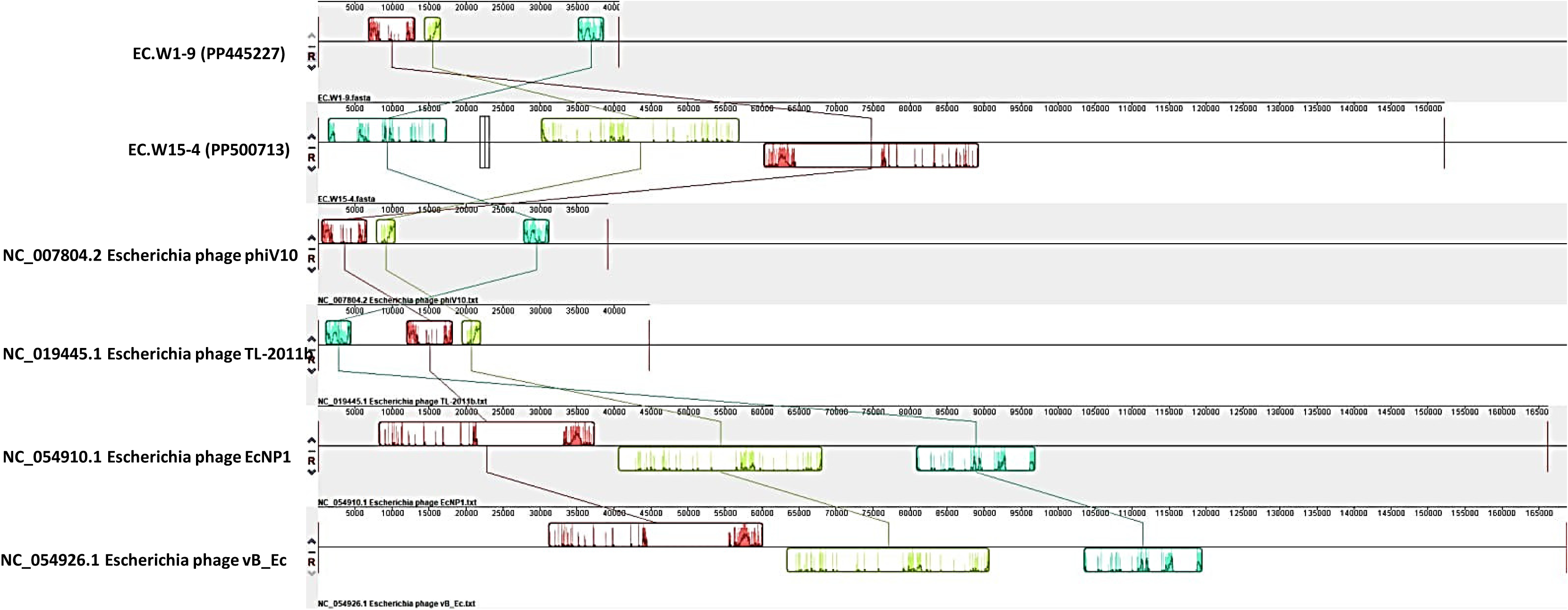
In this figure, this study employed the progressive Mauve algorithm (v2.3.1) to perform multiple alignments of the genomes of 2 novel *E. coli* bacteriophages and 6 closest genomes of *E. coli* bacteriophages. The objective was to investigate the rearrangement patterns and synteny in these genomes. The genomes were laid out horizontally, with homologous segments represented by coloured blocks. The regions outside the blocks lacked homology and were represented by white areas, which were unique to each genome and not aligned

**Table 2.** Genomic information of phage EC.W1-9 and EC.W15-4 *E.coli* Bacteriophages.

| Name of phage | EC.W1-9 | EC.W15-4 |
| --- | --- | --- |
| Accession number | PP445227 | PP500713 |
| Isolation Host | KBN4004 (ST648) | KBN7288 (ST131) |
| Class | *Caudoviricetes* | *Caudoviricetes* |
| Family | *Podoviridae* | *Myoviridae* |
| Genus | *Uetakevirus* | *Tequatrovirus* |
| G+C | 45.8 | 36.1 |
| Genome size (BP) | 40,616 | 152,184 |
| Hypothetical protein | 35 | 80 |
| Crispr | 1 | 1 |
| Phage finder | 9 | 93 |
| Integration/excision | - | 1 |
| Replication/recombination/repair | 2 | 9 |
| stability/transfer/defense | 1 | 2 |
| Alien hunter | 1 | 1 |
| Endolysin | 1 | - |
| Holin | 1 | 1 |
| Endopeptidase | 1 | - |
| Tail fiber/spike protein | 3 | 12 |
| Lysis protein | 1 | 5 |
| Putative peptidase | 1 | - |
| Endonuclease | - | 6 |
| Glutaredoxin | - | 4 |
| Thioredoxin | - | 3 |
| Packaging machinery | 2 | - |
| Head protein | - | 3 |
| Thymidylate synthesis | - | 2 |
| Capsid protein | - | 6 |
| Baseplate protein | - | 1 |
| Genomic similarity | Escherichia phage phiV10 | Escherichia phage vB_EcoM_OE5505 |
| Query Cover /Per. Ident (%) | 66/96.33% | 97/97.94% |

### Comparative evaluation of *in vitro* lytic activity of individual phage’s and their combination

This study compared individual phage and combination to determine their ability to inhibit bacterial growth in ESBL-producing and CREC isolates ((KBN7288 (ST131) KBN4004 (ST648) and KBN6241 (ST410). Analysing bacterial growth curves over 24 h revealed that individual phage and phage combinations demonstrated significant inhibition of all *E. coli* isolate growth in the study. Specifically, from 0 to 10 h, the individual phage and the combination significantly suppressed *E. coli* ATCC25922 and KBN4004 growth. However, during the 6 to 10 h, the combination of EC.W1-9 and EC.W15-4 showed slightly lower inhibition than the individual phage. Notably, after 24 h, the phage combination (EC.W1-9 + EC.W15-4) demonstrated a high level of inhibition against bacterial growth (Fig. 6A-B). Similarly, when evaluating the lytic activity of individual phage and the phage combination against the *E. coli* KBN7288, this study observed no significant changes in growth inhibition within the first 6 h. During the 0 to 10 h, single phage EC.W1-9 provided better lytic activity. However, after 24 h, the combination phage showed better activity against *E. coli* KBN7288 showed growth after 24 h (Fig. 6C). The study also found the combination of phage EC.W1-9 and EC.W15-4 effectively inhibited the growth of *E. coli* KBN 6241, with EC.W1-9 showing a significant inhibition in the first 5 h (Fig. 6D).

**Figure 6:**
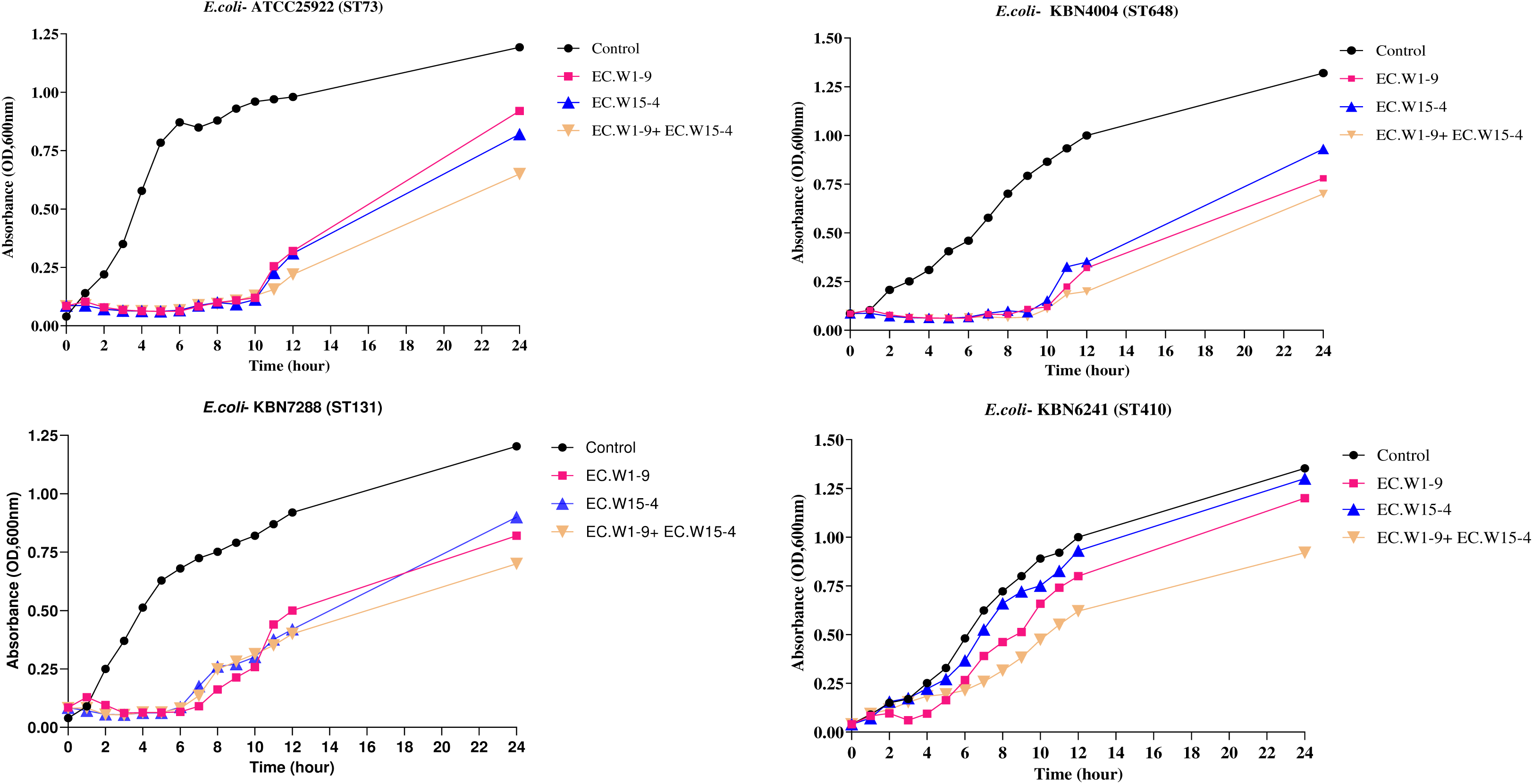
Comparison of the *in vitro* lytic activity of EC.W1-9 and EC.W15-4 phages alone and in combination.

### Assessment of phage-antibiotic synergy (PAS) effect on selected ESBL-producing and CREC isolates

The present study investigated synergistic interactions between phages (EC.W1-9 and EC.W15-4) and antibiotics (Table 3 & S. file 2). When colistin was combined with phage EC.W1-9 against ESBL-producing *E. coli* KBN4004 and ESBL-producing and CREC *E. coli* KBN7288, the MIC values decreased by half (4 to 2 µg/ml and 32 to 16 µg/ml). However, there was no change in MIC values for ESBL-producing and CREC *E. coli* KBN 6241, KBN 6803, and phage and CREC *E. coli* KBN5617. Combining colistin with phage, EC.W15-4 led to a 4-fold (32 to16 µg/ml) reduction in MIC for KBN 7288 and a 2-fold (4 to 2 µg/ml) reduction for KBN4004 and KBN6241, with no change for KBN6803 and KBN5617. Using a phage combination (EC.W1-9 and EC.W15-4) with colistin resulted in significantly lower MIC values than using individual phage. The most significant reductions in the MIC values were observed in KBN7288 (8-fold), followed by KBN4004 and KBN6241 (4-fold each), KBN5617 (2-fold), and KBN6803 (2-fold) (Fig. S2). The research utilized a combination of phage with five antibiotics to target 12 isolates of ESBL-producing and CREC isolates. The results indicated that colistin, meropenem, and tigecycline had a more effective synergistic impact, while amikacin and ciprofloxacin did not show better synergistic effects. In the presence of the colistin alone, only 5 out of 9 isolates had MIC values of ≤ 4 µg/ml. However, 8 out of 9 isolates showed ≤ 4 µg/ml MIC values after phage-colistin synergism. Likewise, in the case of meropenem, only 27.27% of isolates (n=11) had MIC values of ≤ 4 µg/ml. However, after synergistic effects, 54.54% of isolates showed ≤ 4 µg/ml MIC values. Meropenem, in combination with phages, exhibited a notable reduction in MIC values for *E. coli* KBN7288 (1 to 0.25 µg/ml), KBN5617 (16 to 2 µg/ml), KBN 6241(1 to 0.25 µg/ml) and KBN 2048 (16 to 4 µg/ml). Statistical analysis demonstrated that the median MIC value of meropenem alone for all (11) was reduced to 4 µg/ml from 16 µg/ml (4-fold reduction). Similarly, the median MIC value of tigecycline alone (11 ESBL-producing and CREC) was 4 µg/ml, and in combination with the phage cocktail, it was reduced to 1 µg/ml (4-fold reduction). Also, in the case of tigecycline, only 45.45% of isolates (n=11) had MIC values of ≤ 4 µg/ml; however, after synergistic, 90.90% of isolates showed MIC values of ≤ 4 µg/ml. Where a combination of phage and tigecycline better reduces KBN7288 (4 to 0.25 µg/ml), KBN3979 (8 to 1 µg/ml), KBN6241 (4 to 1 µg/ml), KBN6201(4 to 1 µg/ml) and others isolates reduced about 2-fold from the original MIC value of tigecycline. The amikacin and ciprofloxacin provide synergistic effect against ESBL-producing and CREC *E. coli* KBN7288 and reduction MIC (128 to32 µg/ml), Where phage resistant, CREC *E. coli* KBN5617 MIC reduction (16 to 8 and 4 to µg/ml).

**Table 3.**
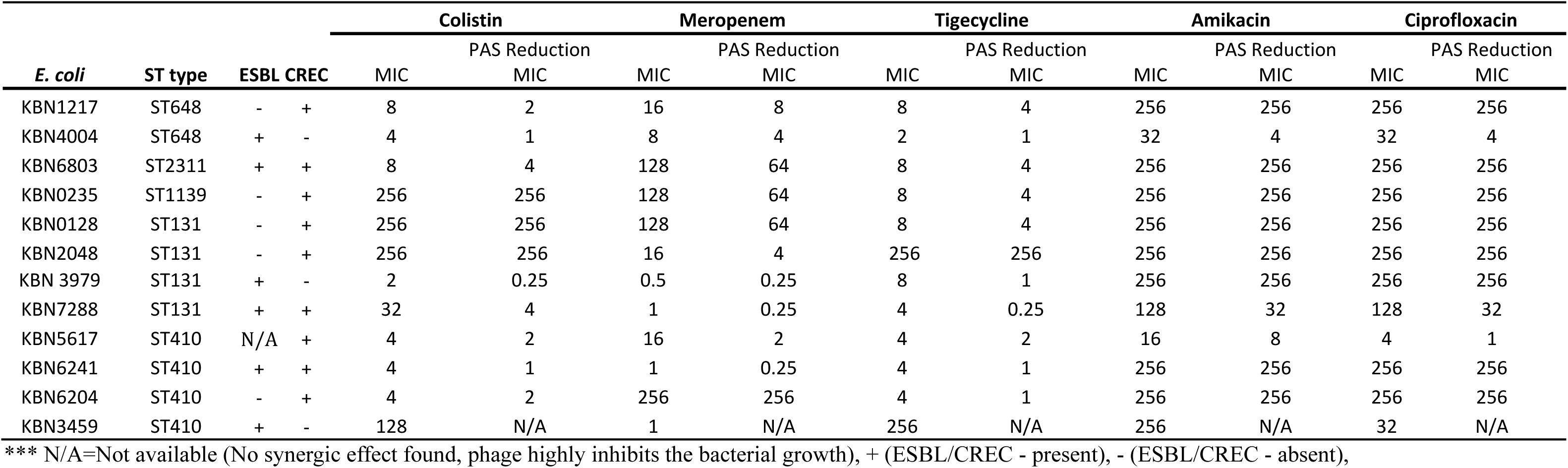
Reduction in MIC values due to PAS among combination phage (EC.W1-9+ EC.W15-4) and antibiotics combination against ESBL-producing *E.coli* and CREC.

### Assessment of the efficacy of phages to treat ESBL-producing and CREC *E. coli* ST 131, ST 648 and ST410 *in vivo*

This study evaluates the efficacy of isolated phages (EC.W1-9 and EC.W15-4) and their combination with antibiotics to treat infections caused by ESBL-producing *E. coli* and CREC in BALB/c mice (Fig. 7). Treatment groups were given phages EC.W1-9, EC.W15-4, antibiotics, and their combinations. The combination of phage (EC.W1-9 + EC.W15-4), and colistin (4 μg/ml) achieved around 100% survival within 7 days against ESBL-producing and CREC *E.coli* KBN7288 (ST131). The combination of phages (EC.W1-9 + EC.W15-4) resulted in approximately 75% survival. Increased survival rates by 50%, 25%, and 50% compared to individual treatments with phage combination and colistin (4 µg/ml), respectively. Single phage treatments resulted in lower survival rates, whereas mice treated with both phage and colistin had the highest survival rates. The untreated positive control group had 100% mortality within 2 days, while the negative control group survived completely. Against ESBL-producing and CREC *E. coli* KBN6241 (ST410), both the combination of phage and the phage-antibiotic combination achieved 100% survival. The positive control group experienced 100% mortality within 3 days. For ESBL-producing *E. coli* KBN4004 (ST648), the phage-antibiotic combination resulted in 100% survival, which was 50% higher than using phage combination alone, which only achieved 50%. When phage and antibiotic treatments were combined, survival were increased over phage combinations alone, with approximately 75% survival against phage-resistant and CREC *E. coli* 5617 (ST410). The study monitored the body weight of mice to indicate health and treatment efficacy. Mice treated with phage and phage-antibiotic combinations showed weight gains, indicating positive therapeutic effects. In the untreated positive control group, there was no weight gain, which aligns with high mortality rates and poor health outcomes (Fig. S3).

**Figure 7.**
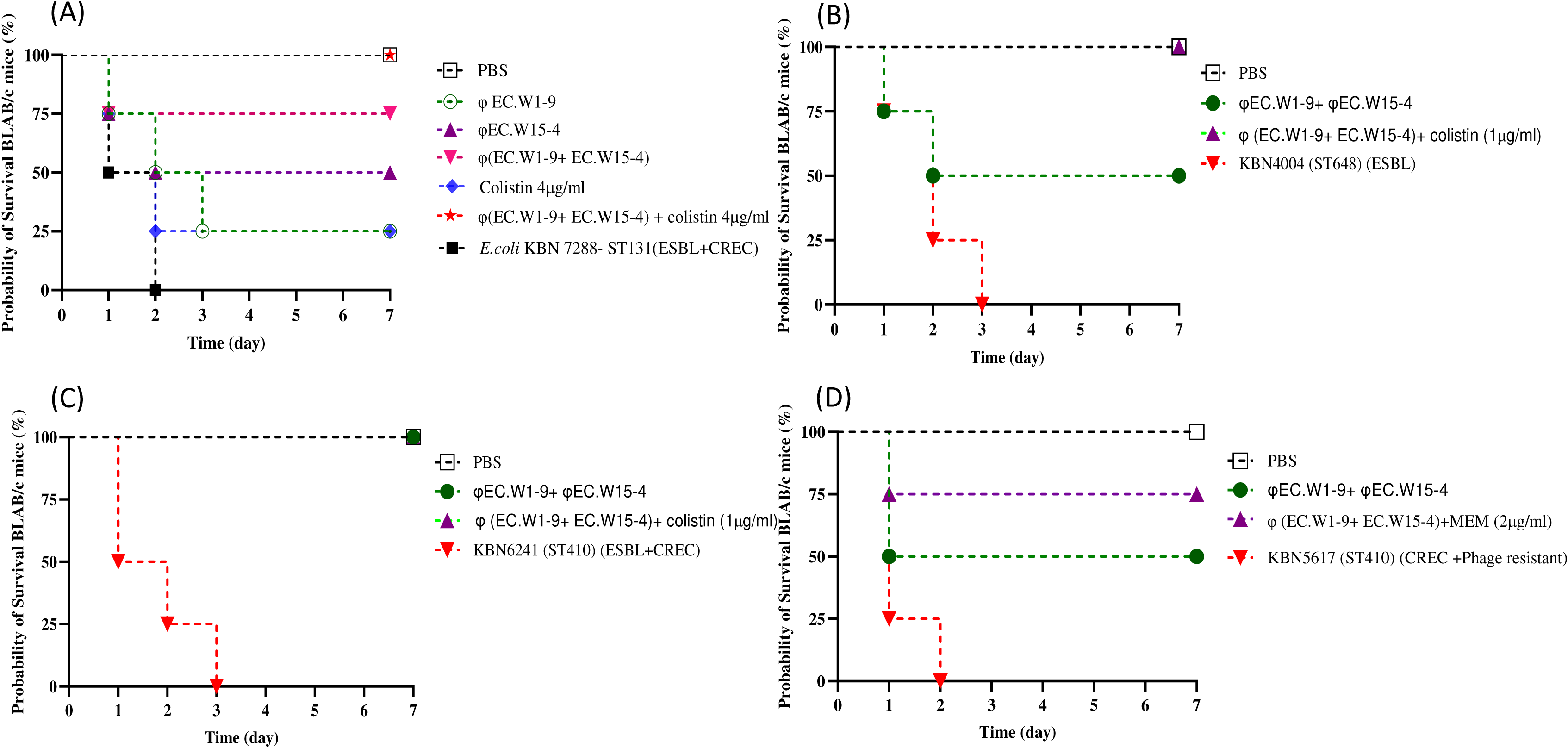
The study examines the toxicity of various ST type of ESBL-producing *E. coli* and CREC in BALB/c mice, assesses survival in phage therapy, and evaluates the efficacy of single phage, phage combinations, and phage-antibiotic combinations (A) Survival curve of treated with phage EC.W1-9 and EC.W15-4, their combination, and the combination of phage’s with colistin (4μg/ml) against ESBL-producing and CREC *E.coli* ST131. (B) Phage EC.W1-9 and EC.W15-4 combination, and with colistin (1μg/ml) against *E. coli* KBN 4004 (ST648). (C) Phage EC.W1-9 and EC.W15-4 combination, and with colistin (1μg/ml) against *E. coli* KBN 6241 (ST410).(D) Phage EC.W1-9 and EC.W15-4 combination, and with meropenem (2μg/ml) against phage-resistant *E. coli* KBN 5617 (ST410).

## DISCUSSION

Antibiotic resistance, specifically ESBL-producing *E. coli* and CREC, poses a significant challenge for global healthcare systems (35). ESBL-producing *E. coli* can degrade multiple antibiotics, making them ineffective for treating infections (11). *E. coli* are resistant to carbapenems, the last resort antibiotics for severe infections (1). To solve this issue, effective strategies must be implemented to maintain antibiotic efficacy and secure treatment alternatives for future generations. Phages show promise in reducing ESBL-producing and CREC *E. coli* (36)(37). The study aimed to isolate and characterize two phages that target 14 distinct sequence types (ST) of ESBL-producing and CREC *E. coli* and determine the efficacy in both *in vitro* and *in vivo*. Following that, this study showed the application of phages as a promising approach for biocontrol ESBL-producing *E. coli* and CREC, such as clones ST131, ST410, ST648, and others. The phages’ morphological features and sequence analysis show strong similarities to the *Podoviridae* and *Myoviridae* families, making them promising candidates for phage-based therapies against *E. coli* infections, as they effectively target and eliminate ESBL-producing *E. coli* and CREC, combating antibiotic resistance (38, 39).

The whole-genome analysis of two *E. coli* phages revealed that they lack virulence, antibiotic resistance, and bacterial toxin-related genes, making them suitable for treating bacterial infections (40). The identification of genes encoding for endolysin and depolymerases in these phages opens up exciting possibilities for biocontrol applications (41). Phage EC.W1-9 contains an endolysin with 94.4% similarity to *E. coli* phages endolysin (<u>YP_512283.1</u>), this enzyme capable of degrading the bacterial cell wall from within, leading to the lysis of the bacterial cell (42). Depolymerases are enzymes that are encoded by phages infecting encapsulated bacteria, typically within open reading frames linked to structural proteins, often found in tail fibers, base plates, head protein and neck regions (43). The analysis of isolated phage genomes revealed the presence of several tail fiber proteins with a high degree of similarity, ranging from 98% to 100%, to those found in other *E. coli* phages (Table S.3). Additionally, the presence of depolymerases in these phage suggests their ability to target and degrade the extracellular polymeric substances that form the biofilm matrix of bacteria (44). The also presence of various proteins such as holin, cell lysis protein, DNA packaging protein, structural protein, DNA replication/transcription/repair proteins, and putative peptidases in phage EC.W1-9 and EC.W15-4 suggests their potential in combating ESBL-producing and CREC infections in clinical or agricultural settings, where biofilms are a significant challenge for traditional antibiotic treatments (45, 46).

Phage therapy must be specific against the target bacterium and preferably have a broad host range to cover the diversity of the target bacterial population (24). In this research, phage EC.W1-9 and EC.15-4 and their combination demonstrated the ability to lysis ten different sequence types (ST) of ESBL and CREC *E. coli* out of the 14 tested ST types. Moreover, when explicitly targeting ESBL-producing and CREC *E. coli*, the combination of phage exhibited lytic activity of 73% and 56.25%, respectively. The single phage EC.W19 and EC.W15-4 showed lytic activities of 55% and 63.33% for ESBL-producing and 37.50% and 50% for CREC. These findings highlight the potential of phage therapy for combating multidrug-resistant bacterial strains. Previous studies have also demonstrated the efficacy of phage cocktails in providing lytic activity against different ST types of ESBL-producing *E. coli* and CREC (47, 48). However, the significant aspect of this study is that the tested phages and their combination exhibited better efficacy against high-risk ESBL-producing and CREC *E. coli* clones ST131, ST410, and ST648, which are commonly found in antimicrobial-resistant *E. coli* infections in Southeast Asia, including South Korea, Japan (49,50). Phage-antibiotic synergy (PAS) refers to sublethal antibiotic concentrations that enhance the release of phages from bacterial cells. The combination of phages and antibiotics shows promise for reducing antibiotic doses and combating antibiotic resistance during treatment (51). In this study, using combination phage with colistin, meropenem, and tigecycline showed better synergistic effects against different ST type of ESBL-producing and CREC *E. coli*. Phage-colistin synergy decreased MIC values to ≤4 µg/ml in 8 out of 9 isolates, compared to 5 out of 9 with colistin alone. Meropenem-phage synergy reduced MIC values significantly, with a median MIC reduction from 16 µg/ml to 4 µg/ml (n=11). Tigecycline-phage synergy also showed a 4-fold reduction, with 90.9% of isolates achieving MIC values of ≤4 µg/ml (n=11). Previous study has also demonstrated that synergistic effects of the phage cocktail with antibiotics was shown by lowering MIC values of antibiotics (52, 53).

To use bacteriophages to treat human bacterial infections, the translation from *in vitro* activity to *in vivo* efficacy is not guaranteed despite a high success rate (54). However, our investigation into the potential of isolated bacteriophages their combination with antibiotics revealed that they could effectively infect targeted different high-risk clone ST type of ESBL-producing and CREC isolates and survival of mice models. The research found that the combination of phage (EC.W1-9 + EC.W15-4) with colistin (4 μg/ml) achieved approximately 100% survival within 7 days against ESBL-producing and CREC KBN7288 (ST131). Additionally, the study revealed that the combination of phages (EC.W1-9 + EC.W15-4) resulted in about 75% survival, marking a significant increase compared to individual treatments. Similarly, phage combination with colistin 1μg/ml increased the survival 100% against ESBL producing *E. coli* KBN4004 (ST 648) and ESBL-producing and CREC *E. coli* KBN 6241 (ST410). Regarding phage resistance and CREC *E. coli* KBN5617 (ST410), combining phage resulted about 50% survival rate, while adding 2μg/ml meropenem increased the survival rate approximately 75%. Interestingly, phages and their combination were resisted against *E. coli* KBN5617. They did not provide lytic activity in the spot test assay. However, the study revealed that the best synergistic effects were observed *in vitro*, and the best survival rates were achieved for the phage combination with antibiotic *in vivo* experiments. Furthermore, it was noted that phage-resistant mutants incurred a significant fitness cost, such as reduced resistance to serum killing and restored antimicrobial susceptibility, as a trade-off for developing phage resistance (15). This finding aligns with the observation that phage cocktails target various structural sites and metabolic activities of a bacterium, reducing the likelihood of simultaneous mutations affecting all receptors (31). Additionally, other studies have shown promising results regarding the use of phage cocktails in treating bacterial infections. For instance, a phage cocktail was utilized to treat *P. aeruginosa* infection in mice, and no phage-resistant bacterial mutants were observed (55). Similarly, phage-resistant mutants are less frequently observed in animal models or during clinical trials, indicating the potential effectiveness of phage therapy (28, 57).

## CONCLUSION

Finally, this research highlights the urgent global health threat of ESBL-producing and CR *E. coli* isolates. Two novel lytic phage, EC.W1-9 and EC.W15-4, have been isolated from hospital sewage water, demonstrating a broad host range and remarkable efficacy in lysing these pathogenic *E.coli* isolates. Their stability under varying pH and temperature conditions and strong lytic activity *in vitro* make them potential for clinical application. Genomic analysis reveals no virulence or drug-resistance genes, making them suitable for therapeutic use. *In vitro*, experiments confirm the synergistic effect of phage with antibiotics against different ESBL-producing and CREC clone ST types. In the injectional mice model, single phage and phage combination with antibiotic administrations have shown significantly improved survival rates compared to individual phage treatments. Based on this comprehensive evaluation, we propose phage, EC.W1-9 and EC.W15-4 are potential solutions for targeting the pandemic ST type clone, including ST131, ST410, ST648, ST7962, ST73, ST2311, ST405, ST1487, ST13003, and ST167. Their emergence provides hope in the face of the escalating threat of ESBL-producing and CREC *E. coli* bacterial infections, marking a crucial step forward in global health research and intervention.

## MATERIALS AND METHODS

### Animals used in *in vivo* experiments

*In vivo* experiments were conducted on BALB/c mice strain from Yeangnam Bio (166 Palgong-ro, Dong-gu Daegu) with 6-week-old female mice receiving sterile food and water. The procedures followed guidelines set by the National Ethics Committee and approved by the Kyungpook National University Animal Care and Use Committee.

### Collection of *E. coli* isolates and culture conditions

ESBL-producing and CREC isolates were collected from the Kyungpook National University Hospital Culture Collection for Pathogens (KNUH-NCCP). The isolates were cultured on 5% sheep blood agar medium at 37°C for 24 h. After that, they were transferred to a brain-heart infusion (BHI) media and incubated at 37°C for 24 h. Finally, the isolates were preserved at -70°C using a 50% glycerol stock for following studies.

### Antimicrobial susceptibility test (AST) and MLST analysis

The study analyzes the antibiotic susceptibility of 60 clinical *E. coli* isolates against 21 antibiotics from 10 antibiotic families. These families consist of extended-spectrum *β* -lactamases, carbapenem, cephalosporin, tetracycline, aminopenicillin, monobactam, fluoroquinolone, penicillin, aminoglycosides, and sulphonamide-trimethoprim. To determine the minimum inhibitory concentration, antibiotic disks (10μg) and culture materials were purchased from Becton, Dickinson, and Company. Type strain *E. coli* ATCC25922 was used as a control. The isolates were tested in triplicate and categorized as resistant, intermediate, and susceptible based on the Clinical & Laboratory Standards Institute (CLSI-2020) guidelines. Moreover, this study also conducted Multi-Locus Sequence Typing (MLST) analysis by the Department of Microbiology at Kyungpook National University. The study aims to understand the antibiotic susceptibility profiles and genetic diversity of *E. coli* isolates based on the guidelines of the Clinical & Laboratory Standards Institute (34).

### Phage isolation, purification, and preparation

Sewage water samples were collected from Kyungpook National University Hospital (KNUH) in Daegu, South Korea. Then the sewage water was centrifuged at 12,000× g for 10 min to remove debris and fine particles. The supernatant was filtered through 0.22 μm pore size membrane filters (Whatman filters, Sigma-Aldrich, St. Louis, USA) to obtain a filtrate for infecting *E. coli* cultures in the early exponential phase (10^8^ CFU mL−1). The infected cultures were incubatated at 30°C (180 rpm) overnight and then kept at 4 °C for 48h. Then, the supernatants were centrifuged at 12,000× g for 10 min and filtered using a 0.22-mm pore membrane. Bacteriophage titers were determined using the double-layered method, purified, and stored at -70°C in a glycerol-supplemented medium (34).

### Determination of lytic activity of phages against different ST type of ESBL-producing and CREC *E. coli*

The spot-tests method was used to determine the host ranges of phages following the methods described by Shukho Kim (2022)(57). For this, 15µL of phage stock (∼10^13^ PFU/mL) was dropped directly on each *E. coli* lawn (n = 60). After this step, the plates were incubated for 24h at 37°C and checked for clear lytic zone. A clear zone indicates the phages lysis of the *E. coli* isolate (+), while the absence of a clear lytic zone indicates the absence of lysis (-).The efficiency of the plating (EOP) assay was also conducted according to the procedure previously described by Mirzaei (58). The EOP of phage was calculated by dividing the average phage titer on the permissive bacteria host over the phage titer on the original host bacteria. Each test was conducted in triplicate.

### Measurement of adsorption rate and burst size of two novel *E. coli* phages

The phage adsorption rate and burst size were determined following the method described by Marzia Rahman (59). *E. coli* were cultured in Brain Heart infusion (BHI) media, then infected with phages at MOI of 0.0001 and incubated at room temperature. Samples were collected and centrifuged at intervals of 0, 1, 2, 5, 10, and 15 min. The supernatants were used for plaque assays to determine unabsorbed phage titers. The phage’s burst size was determined through a single growth curve experiment. Where, *E. coli* cells were harvested and resuspended in a fresh BHI medium, and the phage was added at a MOI of 0.0001 and allowed to adsorb for 30 min at 4°C. Then, the pellet was resuspended in fresh BHI medium and incubated at 37°C. Samples were collected at 5-min intervals for up to 45 min. The samples were diluted and analyzed for phage plaque counting by using the plaque test. Each of the above experiments was independently repeated three times.

### Thermal and pH stability of two novel *E. coli* phages

The thermal and pH stability of the phage were evaluated following the method described by Zhaohui Tang (26). The phage suspension (∼10^8^ PFU/mL) was incubated at different temperatures for 1h. Moreover, pH stability was determined by incubating phage suspension (∼10^8^ PFU/mL) in BHI broth with pH levels ranging from 2 to 10, and phage titers were determined using the double-layer agar method.

### *In vitro* bactericidal activity of two novel *E. coli* phages under different MOIs

The study anlysis by culturing phages with *E. coli* (ATCC25922) in BHI media at different MOIs and incubating them at 24h at 37°C with shaking. The optical density (OD600nm) was measured using a UV-vis spectrophotometer (Molecular Devices, LLC, San Jose, CA, USA) in 96-well plates at 1h intervals for the first 12 and 24 h. A positive control was used bacterial culture without phage, and BHI media was used as a negative control. Each aliquot was tested three times (26).

### Combination effect of bacteriophage and antibiotic targeting ESBL-producing and CREC isolates

Phage antibiotic synergy testing was performed in with LB medium by Carmen Gu Liu et al (60). ESBL-producing and CREC isolates was grown in BHI for 6 h, centrifuged, washed, and re-centrifuged. Then, the pellets were resuspended in the BHI medium and adjusted to ∼1 × 10^9^ CFU/ml (OD600nm = 1). Then, 100 μl was injected into each microtiter plate well containing the phage and antibiotic concentration checkerboard (50 μl for each antimicrobial). The OD600nm was measured every 1 h at 37°C for 24 h by shaking continuously in a Hercuvan Lab Systems thermo shaker incubator (Hercuvan Lab Systems, London).

### Phage genome sequencing and annotation of two novel *E. coli* phages

Phage DNAs were extracted using the phenol-chloroform method described by Džiuginta Jakočiūnė (2018). To conduct next-generation sequencing (NGS), phage genomic DNA was sequenced using an Illumina Miseq platform in San Diego, CA, USA. The sequencing reads were assembled through the Celemics pipeline, based in South Korea (https://btseq.celemics.com/). To compare genome sequences, BLASTn (https://blast.ncbi.nlm.nih.gov/Blast.cgi) was used for alignment. The RAST online website (https://rast.nmpdr.org/rast.cgi) was used to predict open reading frames (ORFs), and the results were cross-checked and corrected using the NCBI database. Gene function maps were created using a lab-developed custom tool and improved with Geneious Prime 2023.2 (https://www.geneious.com/). Phage proteome rectangular tree circular tree was generated using VIPTree (https://www.genome.jp/viptree/). VICTOR (https://ggdc.dsmz.de/submit_victor_job.php) was used to construction of a complete genome phylogenetic tree based on the whole genome sequence of the isolated phages. Phylogenetic trees of large terminase and minor capsid proteins were constructed using the neighbour-joining (NJ) method, and the Bootstrap method was employed to assess the reliability of the phylogenetic trees (https://blast.ncbi.nlm.nih.gov/Blast.cgi). The Mauve algorithm (v2.3.1) was used to visualize the complete genome sequence similarity between the isolated phages and their closest neighbour, *E. coli* phages (https://darlinglab.org/mauve/mauve.html).

### *In vivo* experiments (Injectional bacteraemia model)

Overnight bacterial cultures were grown to the logarithmic phase at 37 °C (250 rpm). The cultures were washed twice in 1X PBS. The number of viable bacteria was determined by plating on BHA agar plate to calculate the colony-forming units (CFU). Then, mice were given 100μl 50% lethal dose (LD_50_) of 10^9^ CFU intraperitoneal injection (IP). After 1h, infected mice received intraperitoneal injections (IP) of purified phage, phage combination, antibiotic and phage antibiotic combination in 100 μl of PBS (61). The infected mice were divided into different treatment groups for analysis. The control group of infected mice that did not receive any treatment. The positive control group was treated with PBS, while the negative control group received both phage and PBS. Daily survival rates were recorded for each group to determine the effectiveness of phage therapy in preventing mortality (62).

## ACKNOWLEDGMENTS

This research was funded by a grant from the Korea Disease Control and Prevention Agency (Grant No. 2022-ER2202-00)

## DATA AVAILABILITY

The complete genome sequences of phages EC.W1-9 and EC.W15-4 have been deposited in GenBank (https://www.ncbi.nlm.nih.gov/) with accession numbers PP445227 and PP500713, respectively/Supplementary material.

## Synergistic effect of phage combination and antimicrobial agent (Colistin) targeting ESBL-producing and CREC *E.coli*

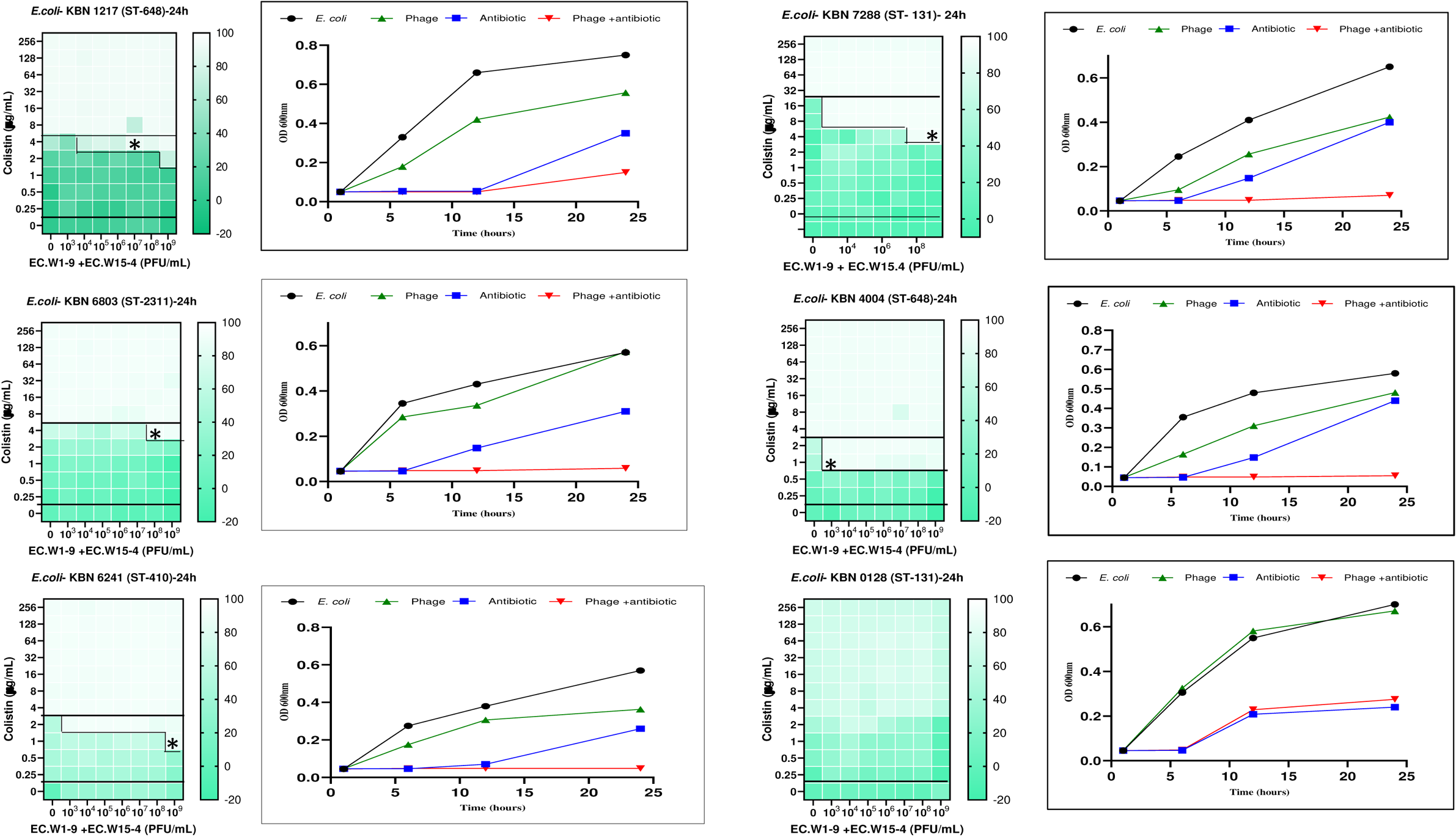
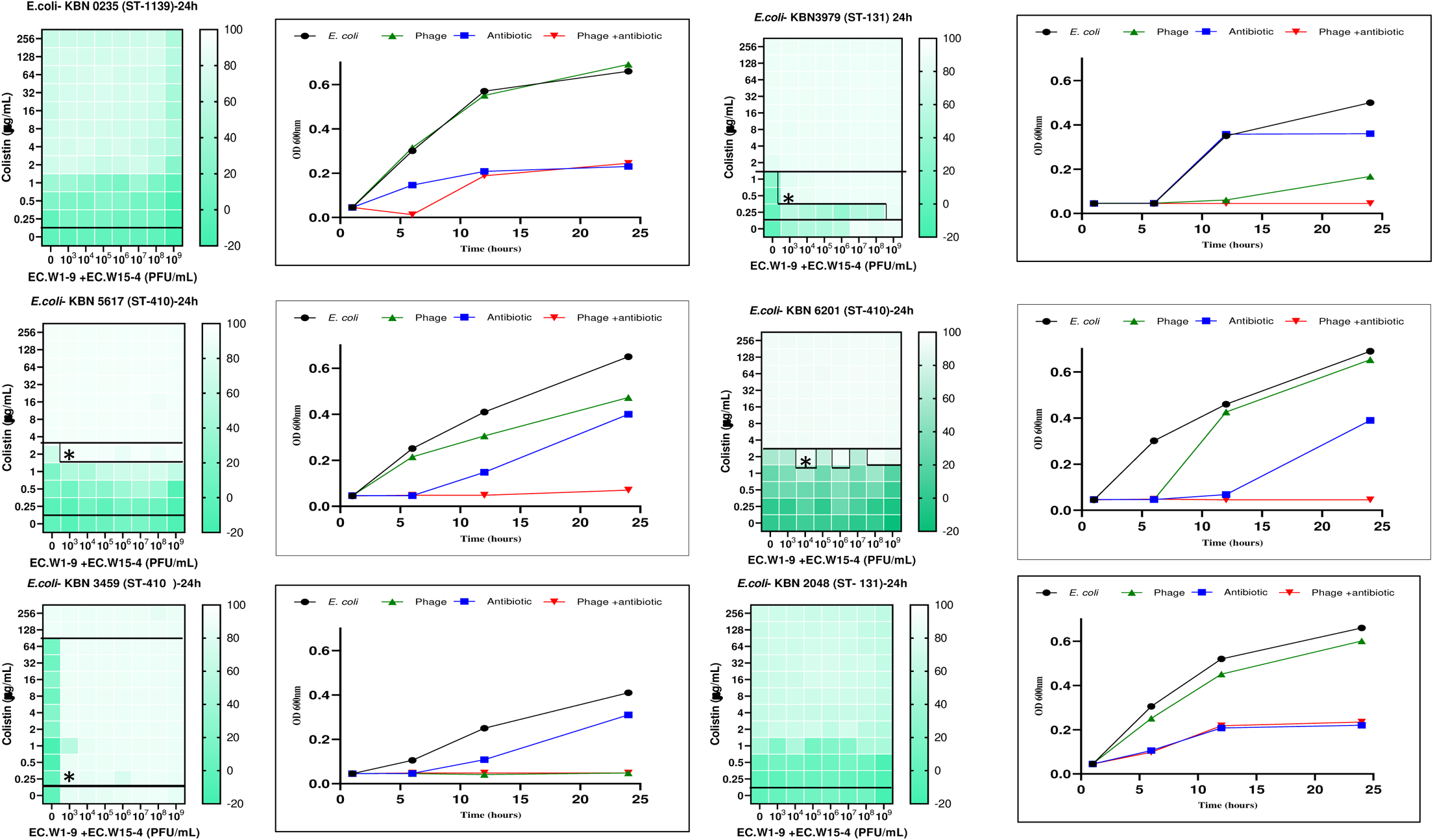

## Synergistic effect of phage combination and antimicrobial agent (Meropenem) targeting ESBL-producing and CREC *E.coli*

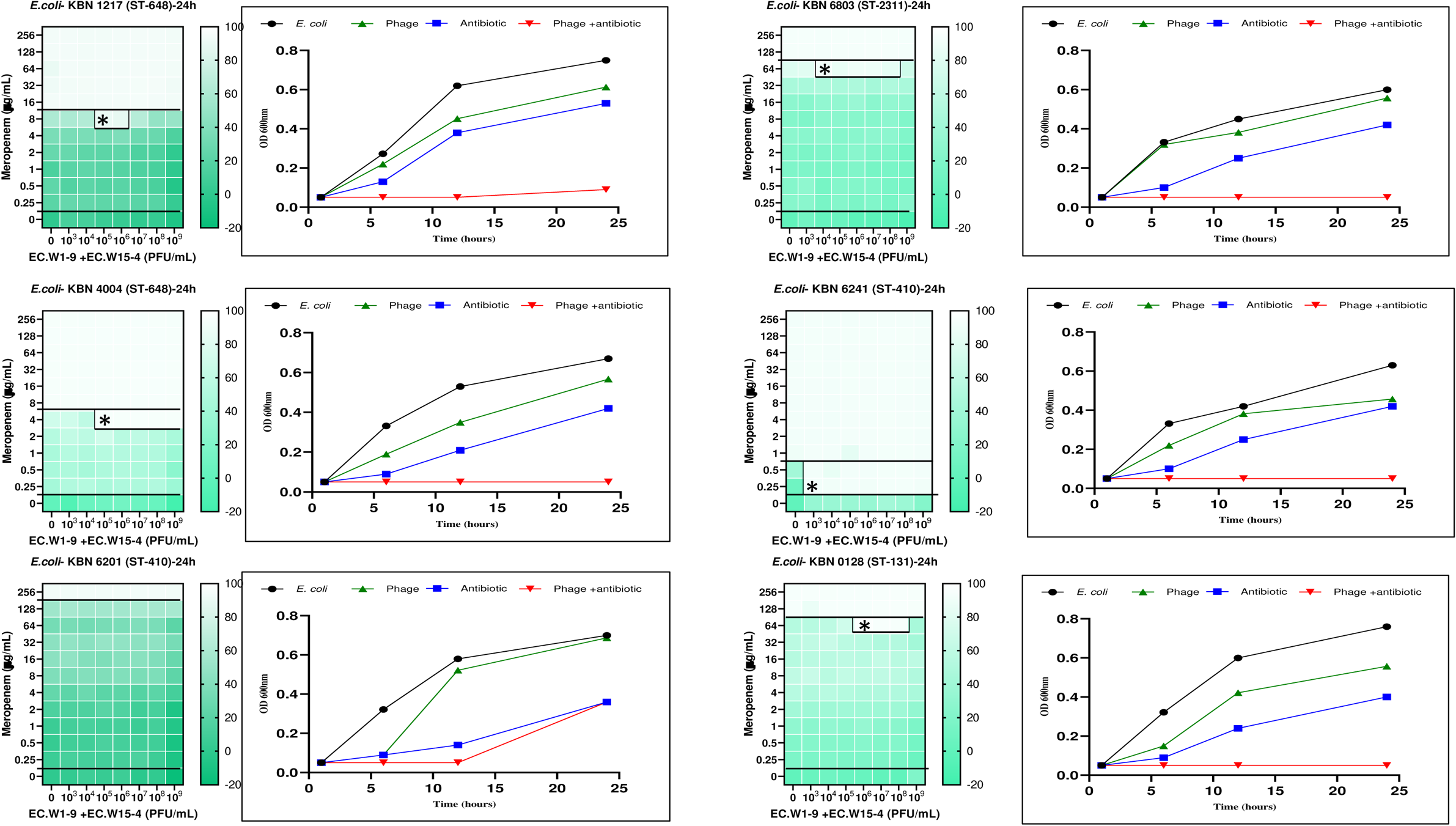
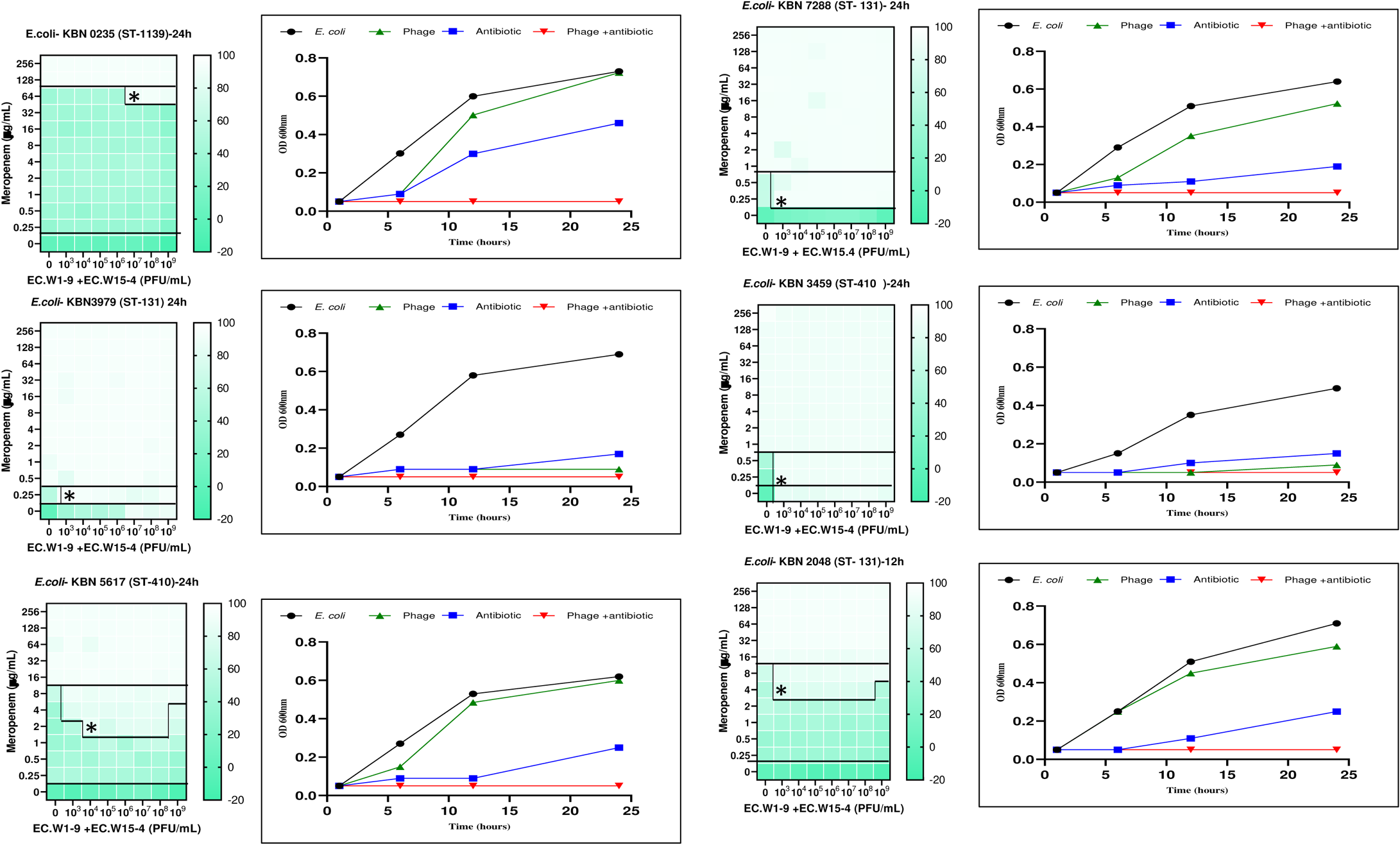

## Synergistic effect of phage combination and antimicrobial agent (Tigecycline) targeting ESBL-producing and CREC *E.coli*

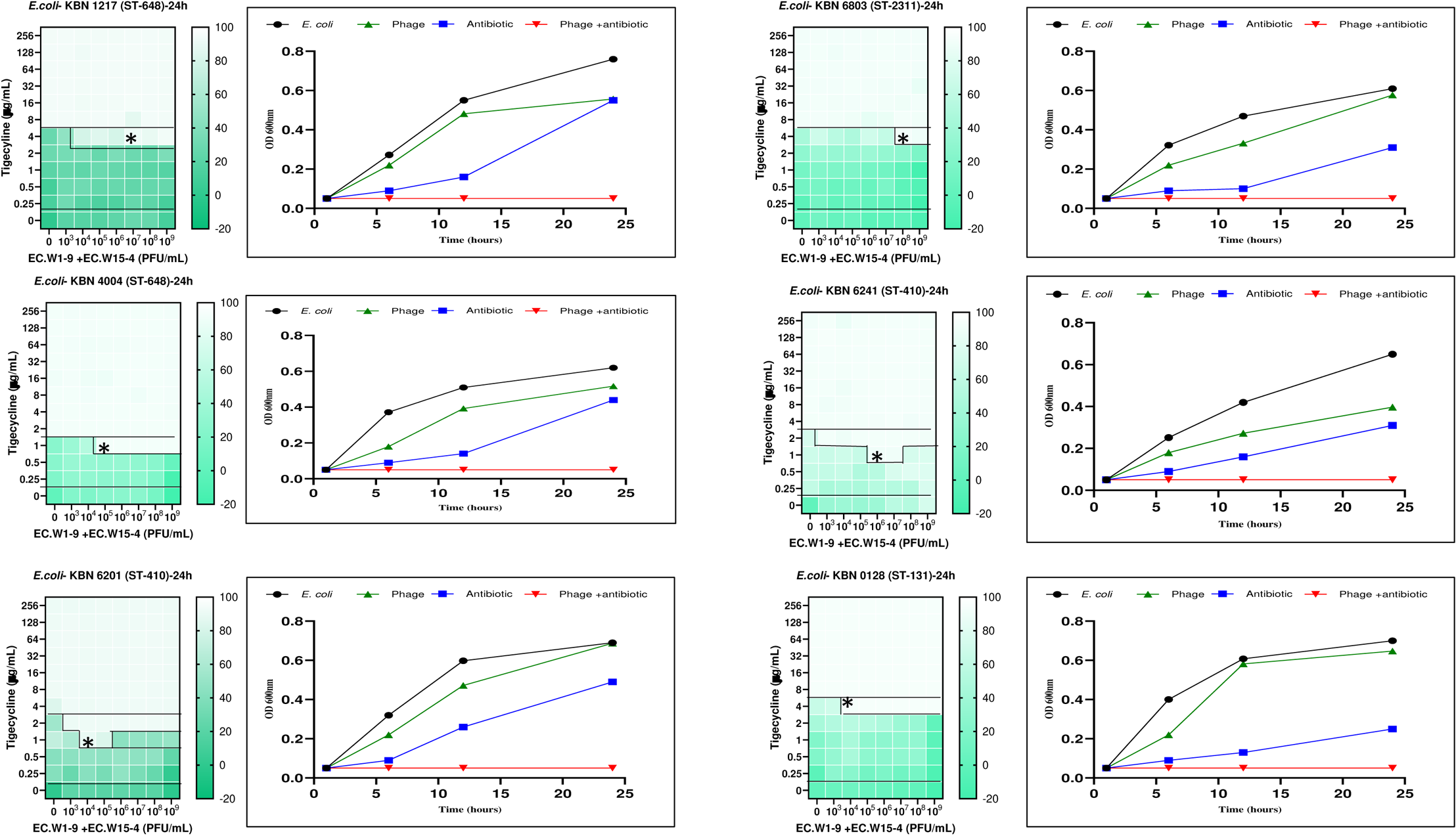
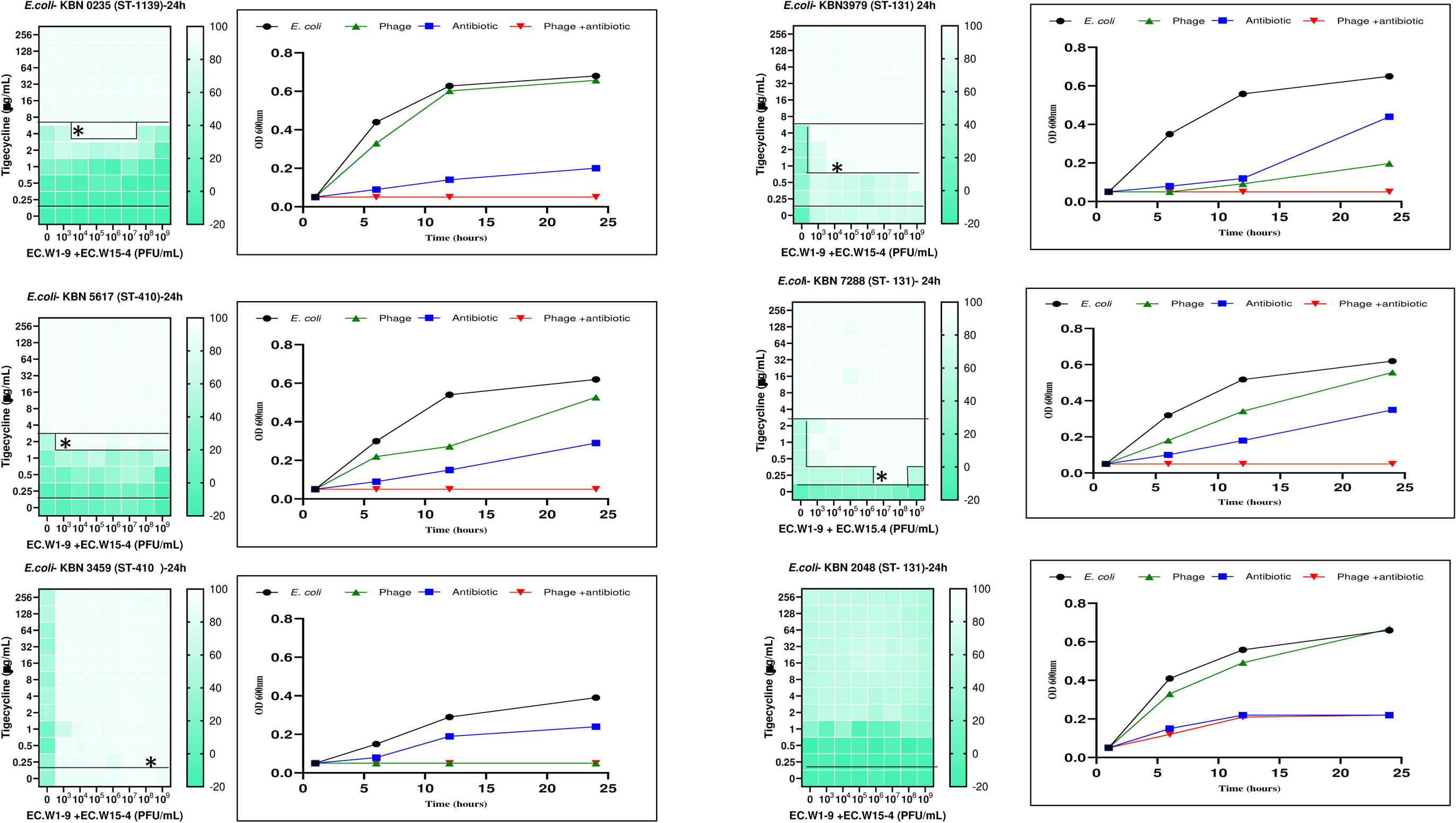

## Synergistic effect of phage combination and antimicrobial agent (Amikacin) targeting ESBL-producing and CREC *E.coli*

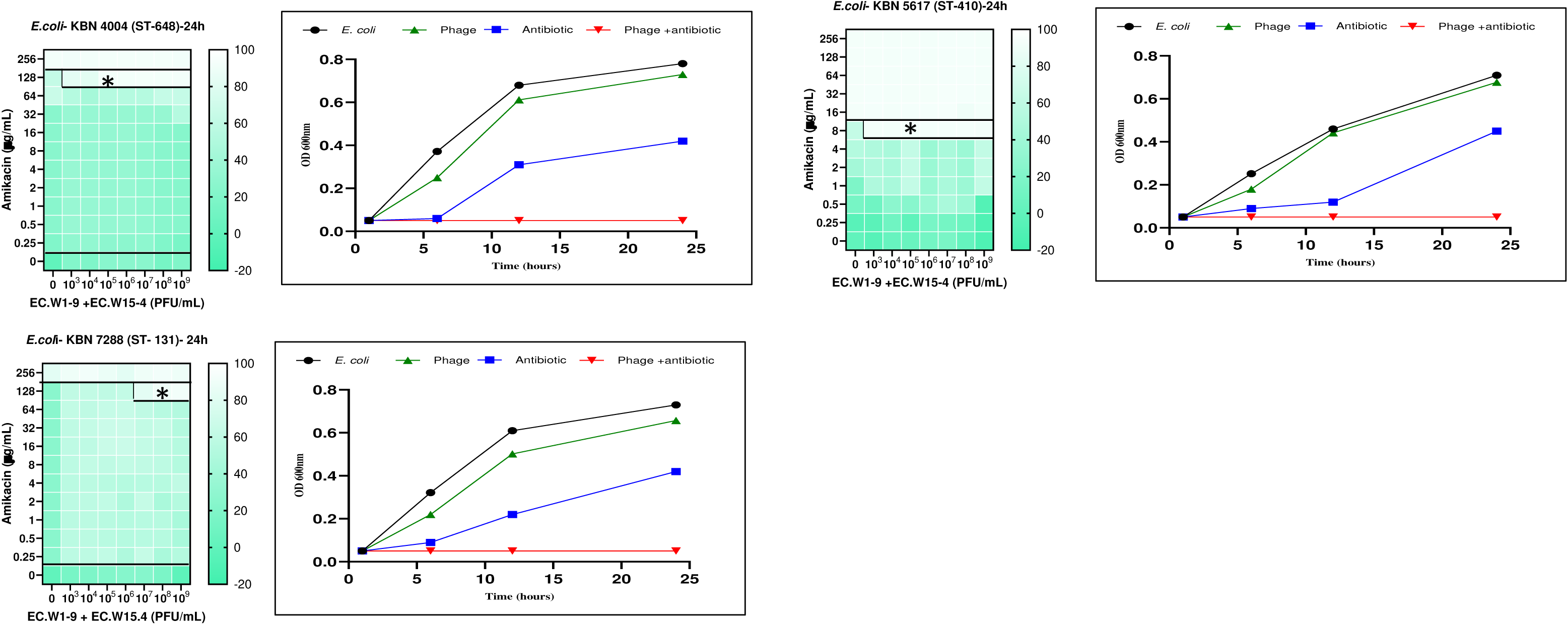

## Synergistic effect of phage combination and antimicrobial agent (ciprofloxacin) targeting ESBL-producing and CREC *E.coli*

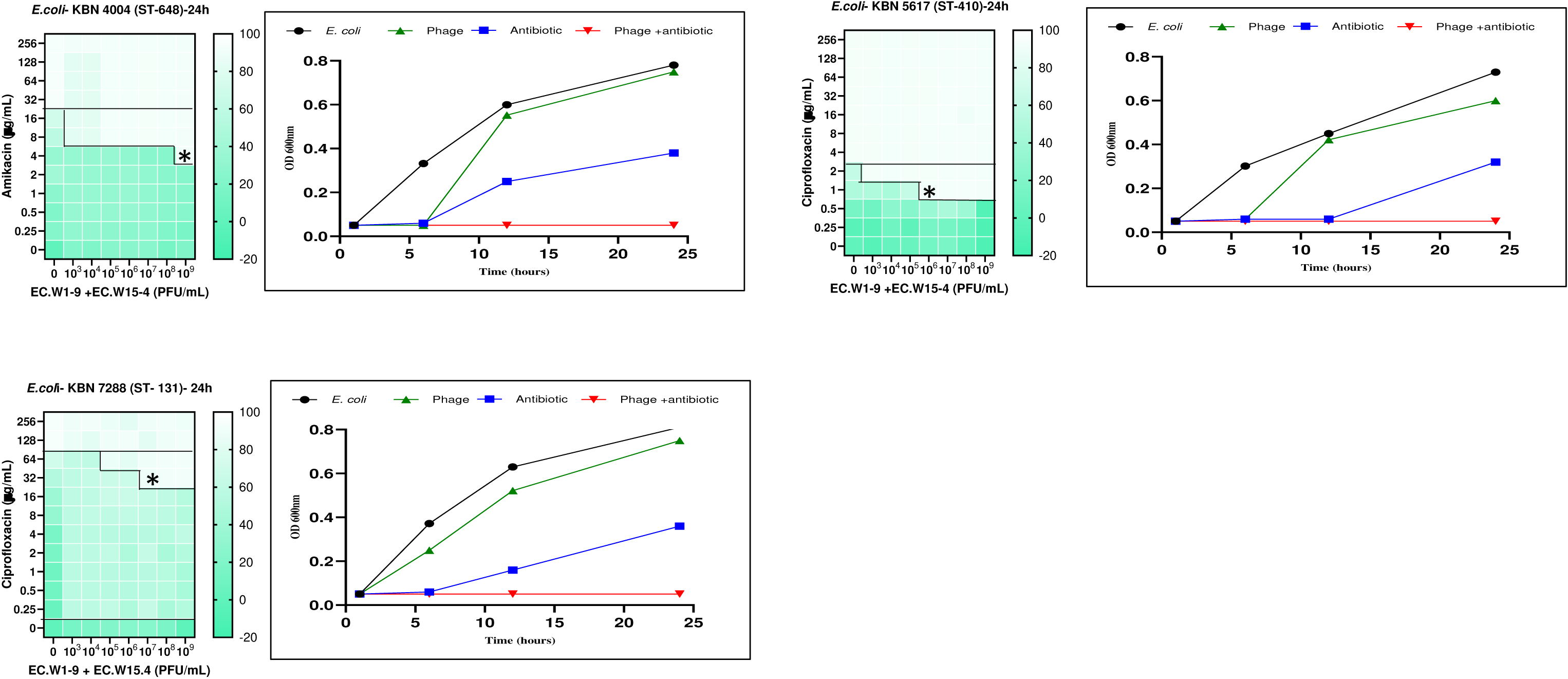

## REFERENCES

1. Huang J, Lv C, Li M, Rahman T, Chang YF, Guo X, Song Z, Zhao Y, Li Q, Ni P, Zhu Y. 2024. Carbapenem-resistant Escherichia coli exhibit diverse spatiotemporal epidemiological characteristics across the globe. Commun Biol 2024 71 7:1–13.

2. Li M, Zhang H, Zhang W, Cao Y, Sun B, Jiang Q, Zhang Y, Liu H, Guo WN, Chang C, Zhou N, Lv C, Guo C, Guo X, Shang J, Huang S, Zhu Y. 2023. One global disseminated 193 kb high-risk hybrid plasmid harboring tet(X4), mcr or blaNDM threatening public health. Sci Total Environ 876:162807.

3. Bajaj P, Singh NS, Virdi JS. 2016. Escherichia coli β-lactamases: What really matters. Front Microbiol 7:184844.

4. Shaikh S, Fatima J, Shakil S, Rizvi SMD, Kamal MA. 2015. Antibiotic resistance and extended spectrum beta-lactamases: Types, epidemiology and treatment. Saudi J Biol Sci 22:90–101.

5. Paterson DL, Bonomo RA. 2005. Extended-spectrum β-lactamases: A clinical update. Clin Microbiol Rev 18:657–686.

6. Quiñones D, Aung MS, Carmona Y, González MK, Pereda N, Hidalgo M, Rivero M, Zallas A, del Campo R, Urushibara N, Kobayashi N. 2020. High Prevalence of CTX-M Type Extended-Spectrum Beta-Lactamase Genes and Detection of NDM-1 Carbapenemase Gene in Extraintestinal Pathogenic Escherichia coli in Cuba. Pathog 2020, Vol 9, Page 65 9:65.

7. Bonnet R. 2004. Growing Group of Extended-Spectrum β-Lactamases: The CTX-M Enzymes. Antimicrob Agents Chemother 48:1–14.

8. Castanheira M, Simner PJ, Bradford PA. 2021. Extended-spectrum β-lactamases: an update on their characteristics, epidemiology and detection. JAC-Antimicrobial Resist 3.

9. Rawat D, Nair D. 2010. Extended-spectrum ß-lactamases in gram negative bacteria. J Glob Infect Dis 2:263.

10. Green SI, Kaelber JT, Li M, Trautner BW, Ramig RF, Maresso AW. 2017. Bacteriophages from ExPEC Reservoirs Kill Pandemic Multidrug-Resistant Strains of Clonal Group ST131 in Animal Models of Bacteremia 10.1038/srep46151.

11. Husna A, Rahman MM, Badruzzaman ATM, Sikder MH, Islam MR, Rahman MT, Alam J, Ashour HM. 2023. Extended-Spectrum β-Lactamases (ESBL): Challenges and Opportunities. Biomed 2023, Vol 11, Page 2937 11:2937.

12. Venkatesan P. 2021. WHO 2020 report on the antibacterial production and development pipeline. The Lancet Microbe 2:e239.

13. Suay-García B, Teresa Pérez-Gracia M. 2019. antibiotics Present and Future of Carbapenem-Resistant Enterobacteriaceae (CRE) Infections 10.3390/antibiotics8030122.

14. Depoorter P, Persoons D, Uyttendaele M, Butaye P, De Zutter L, Dierick K, Herman L, Imberechts H, Van Huffel X, Dewulf J. 2012. Assessment of human exposure to 3rd generation cephalosporin resistant E. coli (CREC) through consumption of broiler meat in Belgium. Int J Food Microbiol 159:30–38.

15. Tian X, Fang R, Wu Q, Zheng X, Zhao Y, Dong G, Wang C, Zhou T, Cao J. 2020. Emergence of a multidrug-resistant ST 27 Escherichia coli co-harboring blaNDM-1, mcr-1, and fosA3 from a patient in China. J Antibiot 2020 739 73:636–641.

16. Gomez-Simmonds A, Annavajhala MK, Wang Z, Macesic N, Hu Y, Giddins MJ, O’Malley A, Toussaint NC, Whittier S, Torres VJ, Uhlemann AC. 2018. Genomic and geographic context for the evolution of high-risk carbapenem-resistant enterobacter cloacae complex clones ST171 and ST78. MBio 9:1–15.

17. Linkevicius M, Bonnin RA, Alm E, Svartstrom O, Apfalter P, Hartl R, Hasman H, Roer L, Raisanen K, Dortet L, Pfennigwerth N, Hans JB, Monnet DL, Kohlenberg A. 2023. Rapid cross-border emergence of NDM-5-producing Escherichia coli in the European Union/European Economic Area, 2012 to June 2022. Eurosurveillance 28:2300209.

18. Zhang R, Li Y, Chen J, Liu C, Sun Q, Shu L, Chen G, Wang Z, Wang S, Li R. 2023. Population genomic analysis reveals the emergence of high-risk carbapenem-resistant Escherichia coli among ICU patients in China. J Infect 86:316–328.

19. Guo CH, Liu YQ, Li Y, Duan XX, Yang TY, Li FY, Zou M, Liu BT. 2023. High prevalence and genomic characteristics of carbapenem-resistant Enterobacteriaceae and colistin-resistant Enterobacteriaceae from large-scale rivers in China. Environ Pollut 331:121869.

20. He W-Y, Lv L-C, Pu W-X, Gao G-L, Zhuang Z-L, Lu Y-Y, Zhuo C, Liu J-H. 2023. Characterization of an International High-Risk Escherichia coli ST410 Clone Coproducing NDM-5 and OXA-181 in a Food Market in China. Microbiol Spectr 11.

21. Brives C, Pourraz J. Phage therapy as a potential solution in the fight against AMR: obstacles and possible futures 10.1057/s41599-020-0478-4.

22. Kim S, Kim SH, Rahman M, Kim J. 2018. Characterization of a Salmonella Enteritidis bacteriophage showing broad lytic activity against Gram-negative enteric bacteria. J Microbiol 56:917–925.

23. Jaglan AB, Anand T, Verma R, Vashisth M, Virmani N, Bera BC, Vaid RK, Tripathi BN. 2022. Tracking the phage trends: A comprehensive review of applications in therapy and food production. Front Microbiol 13:993990.

24. Hibstu Z, Belew H, Akelew Y, Mengist HM. 2022. Phage Therapy: A Different Approach to Fight Bacterial Infections. Biol Targets Ther 16:173–186.

25. Hon K, Liu S, Camens S, Bouras GS, Psaltis AJ, Wormald PJ, Vreugde S. 2022. APTC-EC-2A: A Lytic Phage Targeting Multidrug Resistant E. coli Planktonic Cells and Biofilms. Microorganisms 10:102.

26. Tang Z, Tang N, Wang X, Ren H, Zhang C, Zou L, Han L, Guo L, Liu W. 2023. Characterization of a lytic Escherichia coli phage CE1 and its potential use in therapy against avian pathogenic Escherichia coli infections. Front Microbiol 14:1091442.

27. Koskella B, Brockhurst MA. 2014. Bacteria–phage coevolution as a driver of ecological and evolutionary processes in microbial communities. FEMS Microbiol Rev 38:916–931.

28. Oechslin F. 2018. Resistance Development to Bacteriophages Occurring during Bacteriophage Therapy. Viruses 2018, Vol 10, Page 351 10:351.

29. Wright RCT, Friman VP, Smith MCM, Brockhurst MA. 2019. Resistance evolution against phage combinations depends on the timing and order of exposure. MBio 10.

30. Abedon ST, Danis-Wlodarczyk KM, Wozniak DJ. 2021. Phage cocktail development for bacteriophage therapy: Toward improving spectrum of activity breadth and depth. Pharmaceuticals 14:1019.

31. Malik DJ, Sokolov IJ, Vinner GK, Mancuso F, Cinquerrui S, Vladisavljevic GT, Clokie MRJ, Garton NJ, Stapley AGF, Kirpichnikova A. 2017. Formulation, stabilisation and encapsulation of bacteriophage for phage therapy. Adv Colloid Interface Sci 249:100–133.

32. Forti F, Roach DR, Cafora M, Pasini ME, Horner DS, Fiscarelli E V., Rossitto M, Cariani L, Briani F, Debarbieux L, Ghisotti D. 2018. Design of a broad-range bacteriophage cocktail that reduces pseudomonas aeruginosa biofilms and treats acute infections in two animal models. Antimicrob Agents Chemother 62.

33. Tsai CJY, Loh JMS, Proft T. 2016. Galleria mellonella infection models for the study of bacterial diseases and for antimicrobial drug testing. Virulence 7:214–229.

34. Shamsuzzaman, M., Kim, S. and Kim, J. (2024). Therapeutic Phage Candidates for Targeting Prevalent Sequence Types of Carbapenem-Resistant Escherichia coli https://mc.manuscriptcentral.com/foodborne (Accessed June 5, 2024).

35. Cui L, Zhao J, Lu J. 2015. Molecular characteristics of extended spectrum ß-lactamase and carbapenemase genes carried by carbapenem-resistant Enterobacter cloacae in a Chinese university hospital. Turkish J Med Sci 45:1321–1328.

36. Vitt AR, Sørensen AN, Bojer MS, Bortolaia V, Sørensen MCH, Brøndsted L. 2024. Diverse bacteriophages for biocontrol of ESBL-and AmpC-β-lactamase-producing E. coli. iScience 27:108826.

37. Liang Z, Shi YL, Peng Y, Xu C, Zhang C, Chen Y, Luo XQ, Li QM, Zhao CL, Lei J, Yuan ZQ, Peng YZ, Song BQ, Gong YL. 2023. BL02, a phage against carbapenem- and polymyxin-B resistant Klebsiella pneumoniae, isolated from sewage: A preclinical study. Virus Res 331:199126.

38. Koonjan S, Cooper CJ, Nilsson AS. 2021. Complete genome sequence of vb_ecop_su7, a podoviridae coliphage with the rare c3 morphotype. Microorganisms 9:1576.

39. Wolfram-Schauerte M, Pozhydaieva N, Viering M, Glatter T, Höfer K. 2022. Integrated Omics Reveal Time-Resolved Insights into T4 Phage Infection of E. coli on Proteome and Transcriptome Levels. Viruses 14:2502.

40. Chaudhary N, Singh D, Maurya RK, Mohan B, Mavuduru RS, Taneja N. 2022. Whole genome sequencing and in vitro activity data of Escherichia phage NTEC3 against multidrug-resistant Uropathogenic and extensively drug-resistant Uropathogenic E. coli isolates. Data Br 43:108479.

41. Nakonieczna A, Topolska-Woś A, Łobocka M. 2024. New bacteriophage-derived lysins, LysJ and LysF, with the potential to control Bacillus anthracis. Appl Microbiol Biotechnol 108:1–14.

42. Abdelrahman F, Easwaran M, Daramola OI, Ragab S, Lynch S, Oduselu TJ, Khan FM, Ayobami A, Adnan F, Torrents E, Sanmukh S, El-Shibiny A. 2021. Phage-Encoded Endolysins. Antibiot 2021, Vol 10, Page 124 10:124.

43. Guo Z, Liu M, Zhang D. 2023. Potential of phage depolymerase for the treatment of bacterial biofilms. Virulence 14.

44. Chang C, Yu X, Guo W, Guo C, Guo X, Li Q, Zhu Y. 2022. Bacteriophage-Mediated Control of Biofilm: A Promising New Dawn for the Future. Front Microbiol 13:825828.

45. Wintachai P, Naknaen A, Thammaphet J, Pomwised R, Phaonakrop N, Roytrakul S, Smith DR. 2020. characterization of extended-spectrum-β-lactamase producing Klebsiella pneumoniae phage KP1801 and evaluation of therapeutic efficacy in vitro and in vivo 10:11803.

46. Ali SF, Teh SH, Yang HH, Tsai YC, Chao HJ, Peng SS, Chen SC, Lin LC, Lin NT. 2024. Therapeutic Potential of a Novel Lytic Phage, vB_EclM_ECLFM1, against Carbapenem-Resistant Enterobacter cloacae. Int J Mol Sci 25:854.

47. Uskudar-Guclu A, Yalcin S, Unlu S, Mirza HC, Basustaoglu A. 2023. Evaluation of the Lytic Activity of Various Phage Cocktails Against, ESBL-Producer, Non-Producer and Carbapenem-Resistant Escherichia coli Isolates. Indian J Microbiol 63:208–215.

48. Haines MEK, Hodges FE, Nale JY, Mahony J, van Sinderen D, Kaczorowska J, Alrashid B, Akter M, Brown N, Sauvageau D, Sicheritz-Pontén T, Thanki AM, Millard AD, Galyov EE, Clokie MRJ. 2021. Analysis of Selection Methods to Develop Novel Phage Therapy Cocktails Against Antimicrobial Resistant Clinical Isolates of Bacteria. Front Microbiol 12:613529.

49. Nadimpalli ML, de Lauzanne A, Phe T, Borand L, Jacobs J, Fabre L, Naas T, Le Hello S, Stegger M. 2019. Escherichia coli ST410 among humans and the environment in Southeast Asia. Int J Antimicrob Agents 54:228–232.

50. Tamang MD, Nam HM, Jang GC, Kim SR, Chae MH, Jung SC, Byun JW, Park YH, Lim SK. 2012. Molecular characterization of extended-spectrum-β-lactamase-producing and plasmid-mediated AmpC β-lactamase-producing Escherichia coli isolated from stray dogs in South Korea. Antimicrob Agents Chemother 56:2705–2712.

51. Osman AH, Kotey FCN, Odoom A, Darkwah S, Yeboah RK, Dayie NTKD, Donkor ES. 2023. The Potential of Bacteriophage-Antibiotic Combination Therapy in Treating Infections with Multidrug-Resistant Bacteria. Antibiot 2023, Vol 12, Page 1329 12:1329.

52. Łusiak-Szelachowska M, Międzybrodzki R, Drulis-Kawa Z, Cater K, Knežević P, Winogradow C, Amaro K, Jończyk-Matysiak E, Weber-Dąbrowska B, Rękas J, Górski A. 2022. Bacteriophages and antibiotic interactions in clinical practice: what we have learned so far. J Biomed Sci 29:1–17.

53. Manohar P, Madurantakam Royam M, Loh B, Bozdogan B, Nachimuthu R, Leptihn S. 2022. Synergistic Effects of Phage-Antibiotic Combinations against Citrobacter amalonaticus. ACS Infect Dis 8:59–65.

54. Henry M, Lavigne R, Debarbieux L. 2013. Predicting in vivo efficacy of therapeutic bacteriophages used to treat pulmonary infections. Antimicrob Agents Chemother 57:5961–5968.

55. Yang Y, Shen W, Zhong Q, Chen Q, He X, Baker JL, Xiong K, Jin X, Wang J, Hu F, Le S. 2020. Development of a Bacteriophage Cocktail to Constrain the Emergence of Phage-Resistant Pseudomonas aeruginosa. Front Microbiol 11.

56. Gordillo Altamirano F, Forsyth JH, Patwa R, Kostoulias X, Trim M, Subedi D, Archer SK, Morris FC, Oliveira C, Kielty L, Korneev D, O’Bryan MK, Lithgow TJ, Peleg AY, Barr JJ. 2021. Bacteriophage-resistant Acinetobacter baumannii are resensitized to antimicrobials. Nat Microbiol 2021 62 6:157–161.

57. Kim S, Song H, Jin JS, Lee WJ, Kim J. 2022. Genomic and Phenotypic Characterization of Cutibacterium acnes Bacteriophages Isolated from Acne Patients. Antibiotics 11:1041.

58. Mirzaei MK, Nilsson AS. 2015. Isolation of Phages for Phage Therapy: A Comparison of Spot Tests and Efficiency of Plating Analyses for Determination of Host Range and Efficacy. PLoS One 10:e0118557.

59. Rahman M, Kim S, Kim SM, Seol SY, Kim J. 2011. Characterization of induced Staphylococcus aureus bacteriophage SAP-26 and its anti-biofilm activity with rifampicin. Biofouling 27:1087–1093.

60. Liu CG, Green SI, Min L, Clark JR, Salazar KC, Terwilliger AL, Kaplan HB, Trautner BW, Ramig RF, Maresso AW. 2020. Phage-antibiotic synergy is driven by a unique combination of antibacterial mechanism of action and stoichiometry. MBio 11:1–19.

61. Russo TA, Sharma G, Brown CR, Campagnari AA. 1995. Loss of the O4 antigen moiety from the lipopolysaccharide of an extraintestinal isolate of Escherichia coli has only minor effects on serum sensitivity and virulence in vivo. Infect Immun 63:1263–1269.

62. Pu M, Li Y, Han P, Lin W, Geng R, Qu F, An X, Song L, Tong Y, Zhang S, Cai Z, Fan H. 2022. Genomic characterization of a new phage BUCT541 against Klebsiella pneumoniae K1-ST23 and efficacy assessment in mouse and Galleria mellonella larvae. Front Microbiol 13:950737.

